# Complex developmental and transcriptional dynamics underlie pollinator-driven evolutionary transitions in nectar spur morphology in *Aquilegia* (columbine)

**DOI:** 10.1101/2022.03.24.485692

**Authors:** Molly B. Edwards, Evangeline S. Ballerini, Elena M. Kramer

## Abstract

**Premise:** Determining the developmental programs underlying morphological variation is key to elucidating the evolutionary processes that generated the stunning biodiversity of the angiosperms. Here, we characterize the developmental and transcriptional dynamics of the elaborate petal nectar spur of *Aquilegia* (columbine) in species with contrasting pollination syndromes and spur morphologies.

**Methods:** We collected petal epidermal cell number and length data across four *Aquilegia* species, two with the short, curved nectar spurs of the bee-pollination syndrome, and two with the long, straight spurs of the hummingbird syndrome. We also performed RNA-seq on *A. brevistyla* (bee) and *A. canadensis* (hummingbird) distal and proximal spur compartments at multiple developmental stages. Finally, we intersected these datasets with a previous QTL mapping study on spur length and shape to identify new candidate loci.

**Results:** The differential growth between the proximal and distal surfaces of curved spurs is primarily driven by differential cell division. However, independent transitions to straight spurs in the hummingbird syndrome have evolved by increasing differential cell elongation between spur surfaces. The RNA-seq data reveal these tissues to be transcriptionally distinct, and point to auxin signaling as being involved with the differential cell elongation responsible for the evolution of straight spurs. We identify several promising candidate genes for future study.

**Conclusions:** Our study, taken together with previous work in *Aquilegia,* reveals the complexity of the developmental mechanisms underlying trait variation in this system. The framework we have establish here will lead to exciting future work examining candidate genes and processes involved in the rapid radiation of the genus.

## INTRODUCTION

Angiosperms are the most species-rich and morphologically diverse group in the plant kingdom (Joppa et al., 2011; Soltis et al., 2011), and their origins and rapid evolution are a perpetual source of fascination for biologists (Darwin and Seward, 1903; Stopes, 1912; Regal, 1977; Friedman and Floyd, 2001; Sauquet et al., 2017). Much of angiosperm diversity is apparent in the flower itself — variation in organ identity, size, shape, color, number, fusion, and organization has yielded the seemingly endless floral forms seen in nature. Floral structure is intimately tied to reproductive success, and often adapted to facilitate pollen transfer via animal pollinators; pollinator specialization is associated with rapid diversification in the flowering plants (Armbruster and Muchhala, 2009). Underlying this variation in flower morphology is equally varied developmental genetic pathways, the characterization of which is key to understanding how the angiosperms generated such incredible diversity over the course of their evolutionary history. However, much of our current knowledge of floral organ developmental genetics comes from work done in model systems with relatively simple flowers and few or no pollinator interactions, such as *Arabidopsis* (Bowman et al., 1991; Irish, 2008; Causier et al., 2010; Huang and Irish, 2016). Expanding our investigations into systems with more varied floral forms will reveal new insights into the origins of angiosperm biodiversity.

*Aquilegia* (columbine, Ranunculaceae) is an ideal system for studying the evolution and development of complex floral morphology (Fig. 1A). *Aquilegia* flowers produce an elaborate spur on each of their five petals, which extends downwards from the floral attachment point and ends in a nectary at the tip, while a laminar petal blade extends upwards from the attachment point (Fig. 1B-C). The nectar spur varies dramatically in length, shape, and color depending on what animal pollinates a given *Aquilegia* species: bee-pollinated species tend to have short, curved spurs that are blue-purple in color; hummingbird-pollinated species are typified by red and yellow petals with straight spurs of medium length; and species visited by hawkmoths have extremely elongated, straight spurs (up to 16cm) and have lost floral anthocyanins to enhance crepuscular visibility (Whittall and Hodges, 2007). The genus originated in eastern Asia c. 6.9 million years ago, and the bee-pollination syndrome is ancestral. Upon its migration to North America c. 4.8 million years ago, the lineage encountered new pollinators, including hummingbirds and hawkmoths, and underwent rapid diversification to adapt. There have been two independent transitions from bee-to hummingbird-pollination, during which straightened nectar spurs evolved. This key morphological shift opened up further adaptive potential to hawkmoths, and the multiple subsequent transitions from hummingbird- to hawkmoth- pollination involved dramatic elongation of the nectar spur (Whittall and Hodges, 2007; Fior et al., 2013). A major question in this system (and relevant to angiosperm evolution more broadly) is, What are the developmental and genetic processes that allowed its rapid, pollinator-driven diversification in petal morphology?

**Fig. 1.**
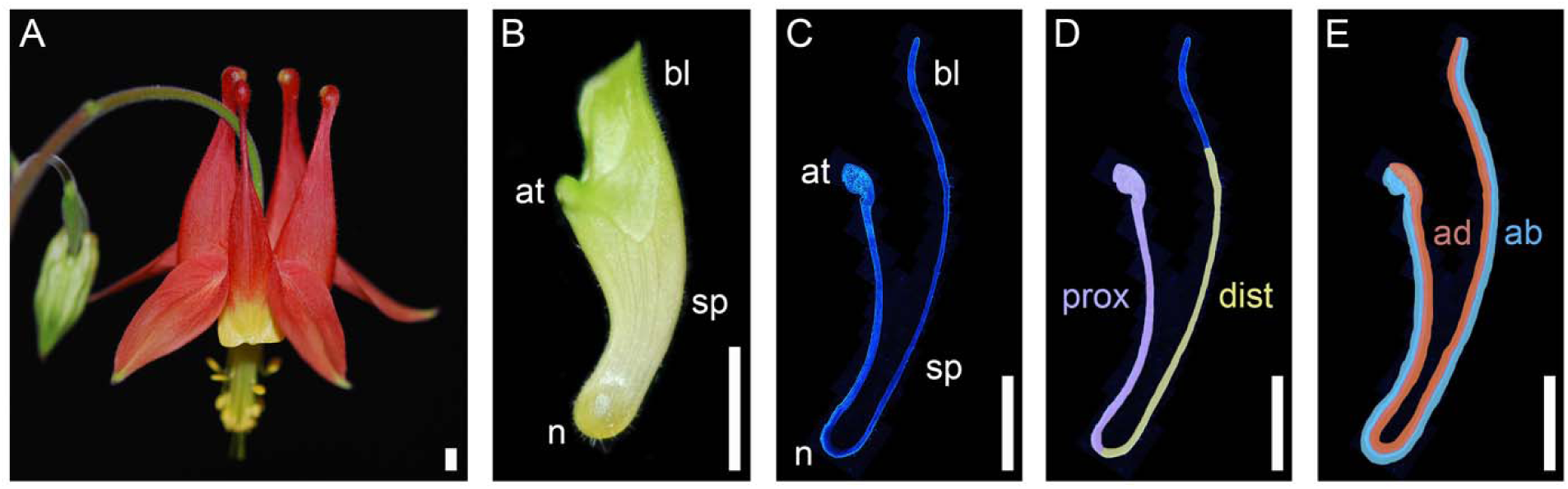
*Aquilegia* petal nectar spurs. **A.** Mature *A. canadensis* flower. **B.** Early *A. canadensis* petal (4mm spur) with attachment point (at), blade (bl), nectary (n), and spur (sp) labeled. **C-E:** Longitudinal section through an early *A. canadensis* petal (6mm spur) stained with calcofluor white. **C.** Same labels as for B. **D.** False-colored to show proximal (prox) and distal (dist) compartments of the spur. **F.** False-colored to show the adaxial (ad) and abaxial (ab) layers of the spur. Scale bar = 2mm.

Previous work on *Aquilegia* nectar spurs revealed two distinct phases of development (Puzey et al., 2012). Phase I is characterized by the concentration of cell divisions at the base of the petal adjacent to the attachment point, resulting in an out-pocketing that forms the nascent spur. Cell divisions cease when the spur is a small fraction of its final length, and remaining spur growth is driven by highly anisotropic cell elongation, which defines phase II of development (Puzey et al., 2012; Yant et al., 2015). Spur length diversity that has been examined to date appears to correlate with the duration of phase II, such that shorter-spurred species exhibit lower petal cell anisotropy than their longer-spurred counterparts (Puzey et al., 2012). However, this work has focused on hummingbird- and hawkmoth-pollinated species, which all have straight spurs; characterizing the development of the curved ancestral state petal and the processes underlying the initial shift to hummingbird pollination is crucial to understanding the rapid diversification of *Aquilegia*.

Creating curvature in a tubular organ requires differential growth between one side of the tube relative to the other, via differential cell division, cell elongation, or both. For *Aquilegia*, we will be referring to these regions of the spur as the proximal and distal compartments, with the proximal being the closest to the attachment point and distal being the furthest (Fig. 1D). The conventional adaxial/abaxial plant lateral organ axes are less relevant for the purposes of this study: the entire epidermal layer inside the *Aquilegia* nectar spur is adaxial, and the entire epidermal layer on the outside of the spur is abaxial (Fig. 1E). Therefore, the proximal and distal compartments of the spur contain both abaxial and adaxial tissues. In the ancestral spur, which is curved towards the attachment point, the distal compartment presumably experiences greater growth than the proximal one. To straighten out during the transition to hummingbird-pollination, growth between the compartments needed to equalize. The nature of these growth patterns is unknown: is cell division, cell expansion, or both the driving factor(s)?

We already have some degree of insight into the genetic basis of these phenomena. We recently conducted a QTL mapping experiment to identify the genetic architecture of both spur length and curvature in a pair of North American sister species, bee-pollinated *A. brevistyla* and hummingbird-pollinated *A. canadensis* (Edwards et al., 2021). This study revealed that spur length has a complex genetic architecture consisting of many loci of small- to moderate-effect, while spur curvature is controlled by a single locus of large effect that explains 46.5% of the curvature variance in the F2s. While this is an exciting preliminary result, a deeper investigation is necessary to identify the developmental and genetic variation responsible for these evolutionarily important phenotypes.

In addition, several genes involved in cell proliferation and expansion in *Aquilegia* nectar spurs have already been characterized, albeit in *A. coerulea* ‘Origami,’ a straight-spurred horticultural hybrid mostly derived from moth-pollinated species. A critical factor in promoting Phase I cell divisions is the C2H2 transcription factor *POPOVICH (AqPOP)* (Ballerini et al., 2020), and the eventual restriction of these divisions to the nascent spur cup is controlled in part by *TEOSINTE BRANCHED/ CYCLOIDEA/ PROLIFERATING CELL FACTOR 4 (AqTCP4)* (Yant et al., 2015). The shift to Phase II anisotropic cell expansion occurs with significant contributions from the auxin and brassinosteroid signaling pathways via *AUXIN RESPONSE FACTOR 6 (AqARF6)* and *AqARF8,* as well as *BES1/ BZR1 HOMOLOG1* (*AqBEH1*), *AqBEH3,* and *AqBEH4* (Zhang et al., 2020; Conway et al., 2021). It is possible that differential expression of these genes in the proximal and distal spur compartments could contribute to variation in spur curvature. We already have some indication that the two compartments are under separate genetic control: silencing of *AqTCP4* resulted in a cellular over-proliferation phenotype specifically in the distal spur compartment (Yant et al., 2015).

With these tantalizing lines of evidence in hand, we set out to more holistically investigate the developmental and transcriptional dynamics of the critical evolutionary shift from curved to straight spurs in *Aquilegia.* In this study, we focus on the proximal-distal spur axis to determine the nature of differential growth and gene expression between compartments in curved spurs, and how those processes shifted to yield straight spurs during the evolution of hummingbird pollination. We again used *A. brevistyla* and *A. canadensis* as our primary study system, as they are the only pair of sister species in the genus with bee- and hummingbird pollination syndromes and are estimated to have diverged c. 3 million years ago (Whittall and Hodges 2007; Fior *et al*. 2013; Fig. 2A). For the developmental component of this work, we examined two additional North American species: bee-pollinated *A. saximontana,* and hummingbird-pollinated *A. formosa,* which evolved the syndrome independently of *A. canadensis* (Whittall and Hodges 2007; Fig. 2A). When synthesized with previous studies in *Aquilegia,* we have uncovered complex mechanisms for spur diversification that suggest a surprising degree of independence between the proximal and distal compartments of the spur. The transcriptomic profiles of these same spur compartments in *A. brevistyla* and *A. canadensis* echo our developmental findings, opening exciting new avenues for exploration, including several promising candidate genes for future investigation.

**Fig. 2.**
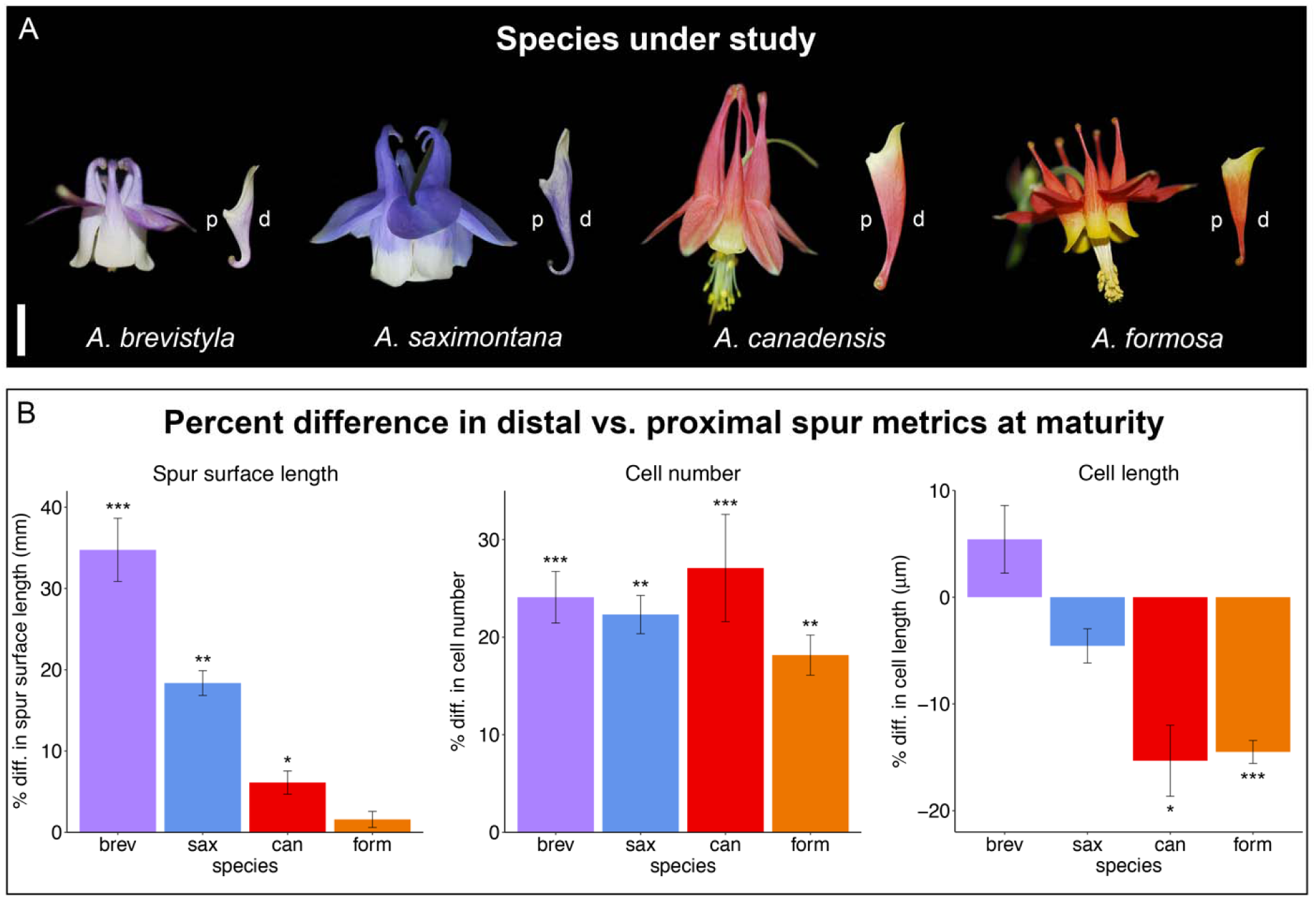
*Aquilegia* nectar spur characterization across species with contrasting pollination syndromes. **A.** Mature flowers and petals of the four North American *Aquilegia* species under study shown to scale: *A. brevistyla* and *A. saximontana* exhibit the ancestral bee-pollination syndrome with blue-purple flowers and short curved nectar spurs, while *A. canadensis* and *A. formosa* represent the two convergently derived instances of hummingbird-pollination in the genus, with red flowers and long straight nectar spurs. Proximal (p) and distal (d) compartments of the spurs are labeled. Scale bar = 1cm. **B.** Percent difference (% diff.) in mature spur metrics (length, cell number, and cell length) of the distal surface vs. the proximal surface of the spur. See Materials and Methods for detailed information on the collection of this data. Error bars represent standard error of mean percent difference across n=6 spurs per species. P-values of t-test to determine significance of difference between proximal and distal measurements are denoted with asterisks: *<0.05, **<0.01, ***<0.001 Abbreviations: brev, *A. brevistyla*; sax, *A. saximontana*; can, *A. canadensis*; form, *A. formosa*.

## MATERIALS AND METHODS

### Plant material and growth conditions

Four *Aquilegia* species were used in this study: *A. brevistyla, A. canadensis, A. formosa,* and *A. saximontana* (Fig. 2A). The *A. brevistyla* plants were descended from a wild population collected near Tucker Lake in Alberta, Canada. *A. canadensis* petal tissue used for developmental analysis was collected from the garden at the Mass Audubon Wildlife Sanctuary in Wellfleet, Massachusetts, and the *A. canadensis* plants used for RNA-seq were descended from a wild population collected in Ithaca, New York. *A. formosa* and *A. saximontana* seeds were purchased from plant-world-seeds.com (Devon, UK). The *A. saximontana* plants from this horticultural supplier were somewhat divergent form the appearance of wild species, so hybridization with other horticultural varieties cannot be ruled out. They still serve their purpose for this study, which was simply to compare *A. brevistyla* to another curved spur. We will refer to them as *A. saximontana* throughout the manuscript.

All species germinated from seed were grown in the Harvard University growth chambers and greenhouses. Seeds were stratified in plug trays on Fafard 3B Soil for 4 weeks at 4 °C, and then transferred to growth chambers set to 16 hour days at 18 °C/13 °C nights to promote germination. After the seedlings outgrew the plug trays, they were moved to 6-inch pots and again to gallon pots. Once they produced 11 mature leaves, they were vernalized at 4 °C for 8 weeks to transition to flowering. Following vernalization, they were transferred to the greenhouse, which kept the same 16-hour-day lighting schedule, but experienced temperatures above 18 °C due to limited cooling ability.

### Developmental measurements and analyses

Petals from the four species were harvested at flower maturity, when approximately half of the anthers had dehisced. They were collected into Farmer’s Fixative (75% ethanol, 25% glacial acetic acid), and fixed for two weeks at 4 °C. Over the course of 12 hours they were gradually dehydrated to 95% ethanol and then stained overnight in 1% Acid Fuchsin (Sigma-Aldrich) in 95% ethanol. Following staining, the petals were rinsed 3x/day in 100% ethanol for several days until the solution no longer turned pink. They were then folded longitudinally, mounted with Cytoseal 60 (Electron Microscopy Sciences, Hatfield, Pennsylvania, USA), sandwiched between a microscope slide and coverslip, and secured with binder clips overnight to ensure they dried flat. They were imaged at the Arnold Arboretum of Harvard University on a Zeiss LSM700 confocal microscope and AxioCam 512 camera; only brightfield observation at 20x magnification was required, but the microscope’s automatic stage enabled the use of the panorama function in the Zen Blue 2.3 software, which allowed seamless movement across the length of the petal to stitch together the dozens of images necessary to capture the entire outline of each spur (Appendix S1). For each spur compartment, only the abaxial epidermal tissue layer was imaged, which will be referred to as the “surface” of the spur.

The microscope images were analyzed in FIJI (Schindelin et al., 2012). On the proximal surface of the spur, cells were counted and measured between the attachment point and the nectary tissue at the tip of the spur. There is no morphological marker comparable to the attachment point on the upper distal surface of the spur, so to consistently determine where to begin distal surface measurements, a line was drawn from the attachment point on the proximal surface to the distal surface of the spur such that it was perpendicular to the distal edge of the spur. Distal surface cells were measured between that point and the nectary tissue at the tip of the spur. The straight-line tool was used to measure the length of each cell along the entire proximal and distal surfaces of the spur, employing several guidelines to maintain consistency in the measurements: first, the length of the cell was always measured parallel to long axis of the spur; second, wherever possible, a continuous file of cells was measured; and third, cells positioned over vascular traces were avoided. The number of cell length measurements served as the cell count for each spur, and the proximal and distal surface lengths were measured using the segmented line tool. Six mature petals were analyzed for each species; cell counts and length measurements were averaged within a proximal or distal surface for a single petal, and then the mean across the six petals was taken for a final average measurement. Percent difference between distal and proximal surface measurements was calculated for each petal using the following formula: Percent difference = ((distal-proximal)/distal) x 100. This value was then averaged across all six petals to determine the mean percent difference for a given spur metric and species.

*A. brevistyla* and *A. canadensis* petals at earlier stages of development were fixed, stained, imaged, and analyzed as for the mature petals, with the modification of higher magnification (40x) and a DIC filter to aid in visualization of the petal cells at the earliest stages. To determine the developmental staging, mature petals from *A. canadensis* and *A. brevistyla* plants were dissected from the flower, laid on their side, and photographed with a Nikon D40X camera with a 60mm lens. Spur length (from attachment point to nectary) was measured from the photographs using the segmented line tool in FIJI (Schindelin et al., 2012) and an average was taken to get a representative measurement of mature spur length in both species. Petal developmental stages were based on percent of mature spur length: 10%, 20%, 30%, 50%, and 70%. All but the 30% stage correspond with the RNA-seq staging (see below). However, petal imaging at the 10% stage was only feasible in *A. canadensis* because the *A. brevistyla* petals were too small at that stage to fold consistently, and imaging at the 70% stage was only necessary for *A. canadensis* to better understand the differential cell elongation dynamics observed in mature petals.

### Tissue collection for RNA-seq

Spur tissue for RNA-seq was dissected and collected according to the following scheme: two species (*A. canadensis* and *A. brevistyla*), four developmental stages (10%, 20%, 50%, 70% of final spur length), two tissue types (proximal and distal), and five biological replicates each, for a total of 80 samples (Fig. 4A, Appendix S2). Petals were placed on an RNAse-free microscope slide resting on ice. The spur was separated from the blade immediately below the attachment point and dissected longitudinally into proximal and distal compartments using a micro scalpel, and then the two halves were separately flash-frozen in liquid nitrogen. Each biological replicate consisted of all five spur-halves from a single flower pooled together; given the time-intensive nature of each petal dissection, the sample collection tubes were suspended in liquid nitrogen so that tissue could be added one-at-a-time rather than performing all five dissections first and then flash-freezing the entire tissue batch, which could have led to RNA degradation. This collection scheme resulted in each biological replicate having a sister sample, which represent the proximal and distal spur compartments from the same flower.

### RNA-seq

Total RNA was extracted from the spur tissue using an RNEasy Plant Mini Kit (Qiagen, Valencia, California, USA), and DNA was removed using the Qiagen on-column DNAse kit during the RNA extraction. RNA quality was assessed on a Nanodrop 8000 spectrophotometer (Thermo Fisher Scientific, Waltham, Massachusetts, USA) and a 2200 TapeStation (Agilent Technologies, Santa Clara, California, USA), and RNA was quantified using a Qubit RNA Broad Range Kit (Thermo Fisher Scientific, Waltham, Massachusetts, USA). Libraries were prepared by the Bauer Core Facility at Harvard University using a KAPA stranded RNA HyperPrep Kit (Roche Sequencing, Wilmington, Massachusetts, USA) and Illumina adapters (Illumina Inc., San Diego, California, USA), and assessed with the 2200 TapeStation and qPCR prior to sequencing. All samples were pooled together and sequenced on 4 lanes of an Illumina NextSeq 500 to generate 38-bp, paired-end reads. Adapters were trimmed using TrimGalore! v0.5.0 (Kreuger, 2019), and kallisto v0.45.1 (Bray et al., 2016) was used to pseudoalign the reads to the *Aquilegia x coerulea ‘*Goldsmith’ v3.1 reference transcriptome (https://phytozome-next.jgi.doe.gov/) and generate abundance calls.

### Differential expression (DE) analysis

The R package edgeR was used for differential expression analyses (Robinson *et al*. 2009; R Core Team 2020). Read normalization was performed with the default calcNormFactors command, which uses the trimmed mean of M-values method, and transcripts that did not have counts per million (cpm) ≥1 for at least five of the 80 samples were filtered out. The z-scores of these normalized filtered reads were calculated using the scale() command in R, and a principal components analysis (PCA) was performed using the prcomp() command. One pair of proximal-distal 20% stage *A. brevistyla* samples clustered with the 10% stage samples, while one pair of 10% stage *A. canadensis* samples clustered with the 20% stage and another 20% stage pair clustered with the 50% stage. These samples were removed from the dataset, and the remaining transcripts were used to estimate dispersions and perform DE analyses.

Pairwise DE between early and late stages within a species and tissue type, or between *A. canadensis* and *A. brevistyla* within a stage and tissue type, was determined using an exact test (exactTest command). For DE analyses between proximal and distal tissue types within a species and stage, a generalized linear model was used with a blocking term for sample pair to account for the fact that each proximal biological replicate is part of a pair with the distal biological replicate that came from the same flower. The quasi-likelihood F-test was used to determine DE genes using the glmQLFTest command.

### Gene Ontogeny (GO) term analysis

Gene Ontogeny (GO) term analysis of gene lists was performed using agriGO v2 (http://systemsbiology.cau.edu.cn/agriGOv2/) (Du et al., 2010; Tian et al., 2017). All expressed genes in the RNA-seq dataset were used as background against which to test for GO enrichment, and all transcript names were converted to *Arabidopsis* locus identifiers based on the *Aquilegia x coerulea ‘*Goldsmith’ v3.1 genome annotation. A Fisher’s exact test and Benjamini-Yekutieli method were used to determine enrichment and adjust for multiple tests, respectively, and the minimum number of mapping entries was set to five. GO terms with an FDR value ≤ 0.01 were considered significantly enriched. Selected GO terms were visualized for figures using code adapted from Martinez *et al*. 2021.

### Weighted gene correlation network analysis

Weighted gene correlation network analysis (WGCNA) was performed on all normalized expressed genes in our dataset using the R package WGCNA v1.70 and methods supplied in the package tutorial (Langfelder and Horvath, 2008). Adjacency was calculated using a soft-thresholding power of 9, the lowest power at which the scale-free topology fit index reached 0.75. The topological overlap matrix was constructed using the default deepSplit value (2), a minModuleSize of 20, and a mergeCutHeight of 0.25. Correlations between the resulting module eigengenes (the first principal component of the expression matrix of a given module) and the sample traits (tissue type, stage, and species) were calculated, and the heat map representing the eigengene-trait relationships was generated using the labeledHeatMap command.

## RESULTS

### Spur development across species with contrasting pollination syndromes

We quantified spur length, cell number, and cell length of the abaxial epidermis of the proximal and distal compartments of mature spurs (which we will refer to as the spur “surface”) across four *Aquilegia* species: *A. brevistyla* and *A. saximontana* with curved spurs, and *A. canadensis* and *A. formosa,* which convergently evolved straight spurs (Fig. 2A). For the following results that compare proximal and distal values for a given spur metric, we are reporting the mean percent difference between those values ± the standard error (Fig. 2B), followed by the P-value of the T-test to indicate whether the mean proximal and distal values themselves are significantly different (Fig. 2B, Appendix S1). The distal surfaces of *A. brevistyla* and *A. saximontana* spurs were 34.8 ± 3.9% (*P* < 0.001) and 18.4 ± 1.5% (*P* < 0.01) longer than the proximal surfaces, respectively, while the distal surface of *A. canadensis* spurs was 6.1 ± 1.4% (*P* < 0.05) longer; however, there was no significant difference in spur surface length in *A. formosa* (P = 0.50; Fig. 2B; Appendix S1). *A. brevistyla* has the shortest spurs of all four species, with an average distal surface length of 11.8 ± 0.2 cm, and *A. canadensis* has the longest spurs, with an average distal surface length of 23.7 ± 0.2 cm (Appendix S1). The distal surfaces of all species’ spurs had significantly more cells than the proximal surfaces, ranging from 18.2 ± 2.1% (*P* < 0.01) more in *A. formosa* to 27.1 ± 5.5% (*P* < 0.001) more in *A. canadensis* (Fig. 2B). Overall, *A. brevistyla* had the lowest cell count of the four species, with 138.7 ± 2.2 cells on the distal surface, while *A. saximontana* had the most with 345.2 ± 13.9 cells on the distal surface (Appendix S1). There was no significant difference in cell length between distal and proximal surfaces of the species with curved spurs (*P* > 0.05), while distal cells were 15.3 ± 3.3% (*P* < 0.05) and 14.5 ± 1.1% (*P* < 0.001) shorter than proximal cells in *A. canadensis* and *A. formosa,* respectively. *A. brevistyla* had the longest distal cells in the study (79.8 ± 1.1 μm), while *A. formosa* had the shortest (54.6 ± 1 μm; Appendix S1). In summary, the curved-spurred species we analyzed appear to generate curvature by producing more cells on the distal surface, while the straight-spurred species achieve their shape through differential cell elongation without dramatically altering the difference in cell number between spur compartments.

To see how these patterns establish themselves through development, we counted and measured spur cells at multiple developmental stages of *A. brevistyla* and *A. canadensis* petals (Fig. 3). Proximal surface cell divisions in *A. brevistyla* spurs cease when the spur is at 30% of its final length, but in the distal surface they cease at 50% of final length. In contrast, they appear to level off at 20-30% of final length in both surfaces in *A. canadensis.* In *A. brevistyla,* cell length increases at a steady rate on both surfaces of the spur throughout development to achieve similar cell lengths at maturity. In *A. canadensis,* the rate of cell elongation increases on the proximal surface of the spur starting at the 30% stage, resulting in proximal cells that are significantly longer than distal cells at maturity.

**Fig. 3.**
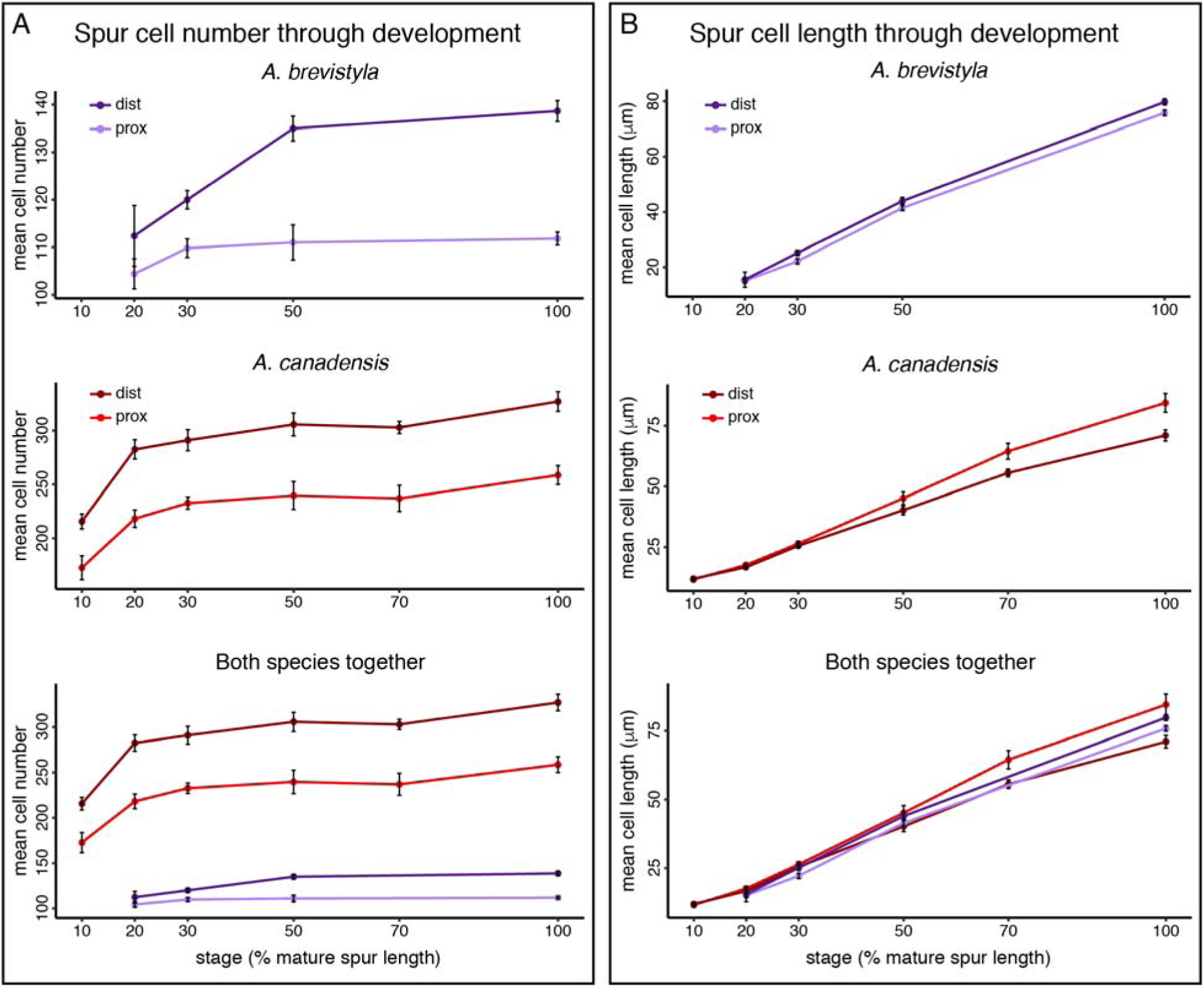
Nectar spur cell characterization through development in *A. brevistyla* and *A. canadensis.* **A.** Spur cell number in proximal and distal surfaces. *A. brevistyla* and *A. canadensis* data are shown separately to better visualize patterns among stages and tissue types, and then again at bottom on the same plot to better compare between species. **B.** Same as A., but for spur cell length.

### RNA-seq and sample clustering

To explore how gene expression varies within and between spurs with these contrasting developmental patterns, we performed RNA-seq on the proximal and distal spur compartments of *A. canadensis* and *A. brevistyla* at four developmental stages (10%, 20%, 50%, and 70%; Fig. 4A, Appendix S2). Five biological replicates of each tissue type were collected, for a total of 80 samples collected from 40 flowers. A total of 1.99 billion raw reads were generated via 38-bp paired-end Illumina NextSeq 500 sequencing, ranging from 19.4 to 31.6 million reads per sample for an average of 24.9 million reads per sample (Appendix S3).

**Fig. 4.**
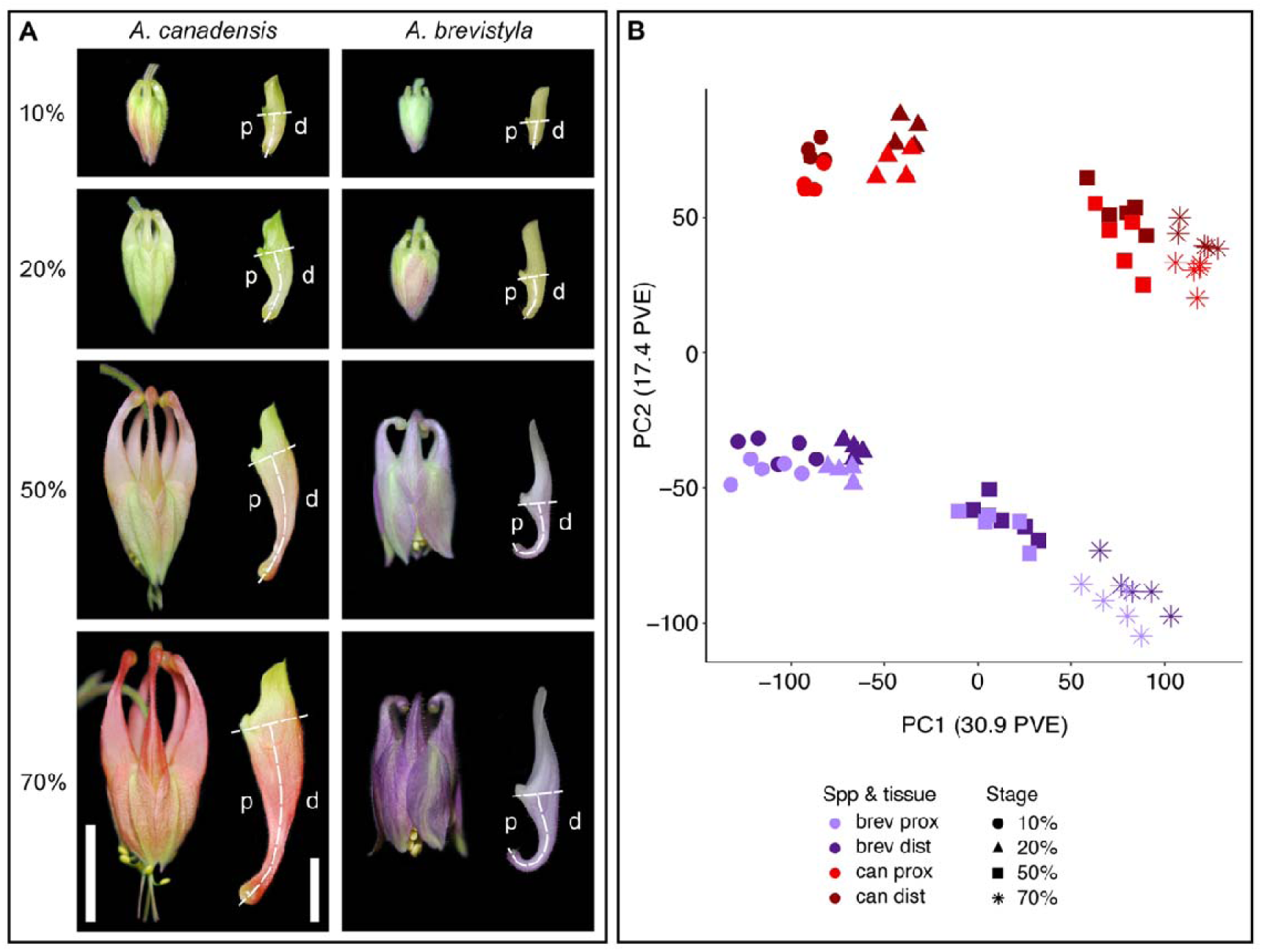
RNAseq experimental design and principal component analysis. **A.** RNAseq sampling scheme. Petals were collected from *A. canadensis* and *A. brevistyla* flowers at four developmental stages based on percent of mature spur length (10%, 20%, 50%, and 70%). For each petal, the blade was removed and the spur was dissected into proximal (p) and distal (d) compartments (dashed lines), which were flash-frozen separately (images of dissected petals are in **Fig. S2)**. Each biological replicate consists of all five spur compartments from a single flower. 80 samples representing 40 flowers were collected in total (2 species, 4 stages, 2 compartments, 5 biological replicates). Flower bud scale bar = 1cm, petal scale bar = 500μm. **B.** The top two principal components (PCs) of the PC analysis on the final normalized RNA-seq dataset. Samples are color- and shape-coded by species (spp), developmental stage, and tissue type. Abbreviations: brev, *A. brevistyla*; can, *A. canadensis*; prox, proximal; dist, distal; PVE, percent variance explained.

Our PCA of the normalized reads shows that the samples generally cluster well together by developmental stage and species (Fig. 4B). PC1 and PC2 together explain 30.9% and 17.4% of the overall variance, respectively. The developmental stages appear to be the main variable affecting PC1, with the earlier two stages and later two stages separated by a large gap on the PC1 axis. PC2 largely captures the differences between species, however there is also a slight separation of proximal and distal samples along that axis.

### Differential expression analysis between earliest and latest developmental stages

We began by comparing gene expression between the earliest and latest developmental stage within each species and tissue type to assess how gene expression changes through time in the developing spur (Fig. 5A). We found a large number of significantly (FDR < 0.05) differentially expressed (DE) genes, ranging from 11,052 in *A. canadensis* proximal tissue to 14,071 in *A. brevistyla* proximal tissue (Appendix S4). *A. brevistyla* had more DE genes overall between the earliest and latest stages than *A. canadensis,* and the earliest stage in all species and tissues had more upregulated genes than the latest stage. The logFC of all significantly DE genes across species and tissue types ranged from -13.7-18.1 (Fig. 5B).

**Fig. 5.**
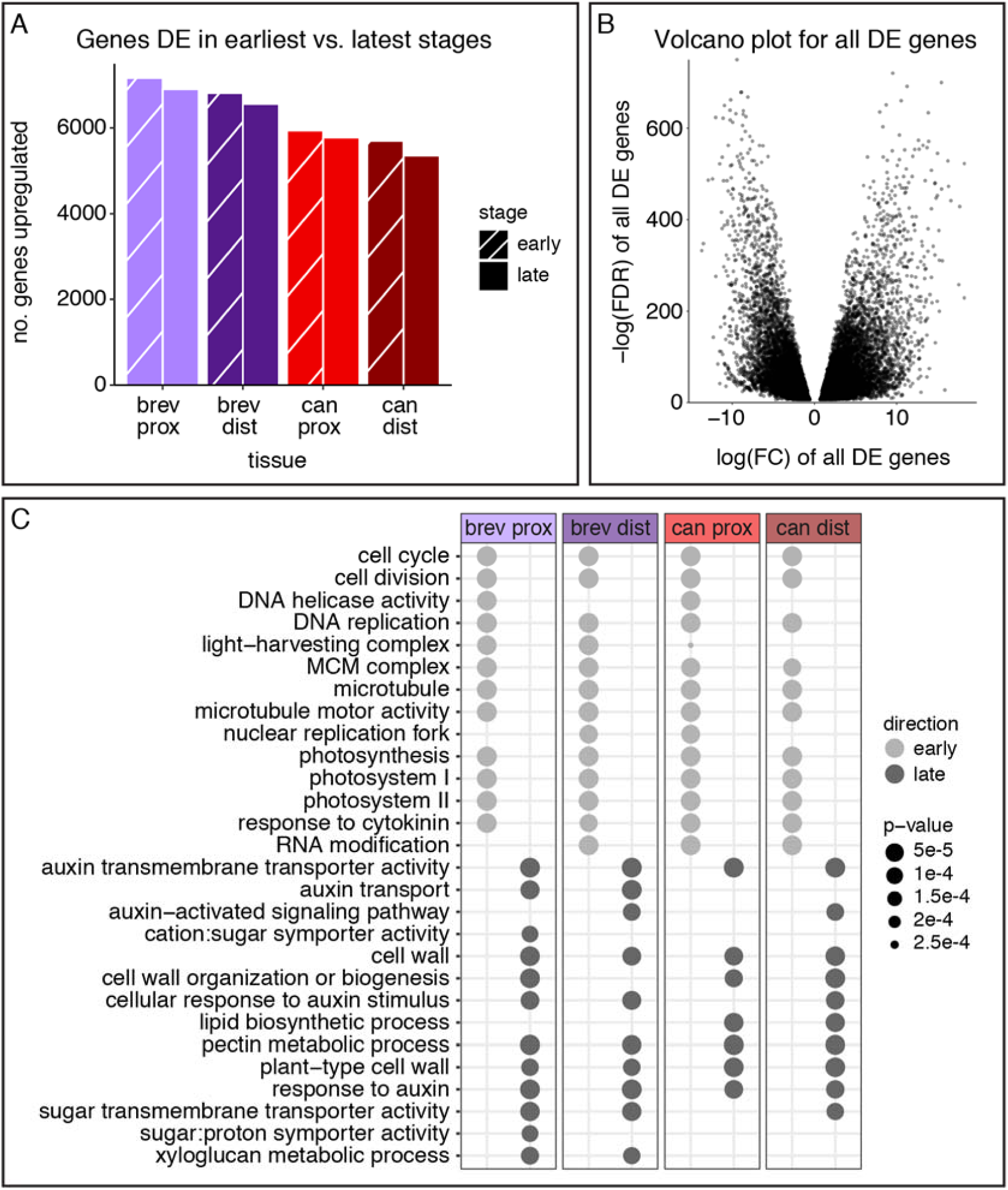
Early vs. late developmental stage DE analysis. **A.** Number of genes DE between earliest (10%) and latest (70%) stages within a species and tissue type. **B.** Volcano plot of all significantly DE genes showing the logFC and -logFDR of each transcript. Points with a positive logFC represent genes upregulated in late tissues, points with a negative logFC represent genes upregulated in early tissues. **C.** Selected GO terms enriched in early and late tissues. Abbreviations: brev, *A. brevistyla*; can, *A. canadensis*; prox, proximal; dist, distal.

A GO term analysis of these DE genes revealed that those upregulated in the earliest stage were enriched for GO terms related to cell division, photosynthesis, and cytokinin response (Fig. 5C, Appendix S5). Genes upregulated in latest stage were enriched for GO terms involved with cell elongation, cell wall synthesis, auxin response, and sugar transport. These GO terms were shared across species and tissue types.

Among the top DE hits in the earliest stage across both species and tissue types was the *Aquilegia* homolog of *AINTEGUMENTA-LIKE 5 (AIL5),* which contributes to petal initiation and growth in *Arabidopsis* (Krizek, 2015). Top DE hits in the latest stage across all sample types included an *Aquilegia* transcript annotated as *XYLOGLUCAN ENDOTRANSGLUCOSYLATE/HYDROLASE 33 (XTH33)*, which is involved in cell wall synthesis and modification in *Arabidopsis* (De Caroli et al., 2021), as well as *PIN-FORMED 5 (PIN5)*, which is localized in the endoplasmic reticulum and regulates intracellular auxin homeostasis in *Arabidopsis* (Mravec et al., 2009).

### Differential expression analysis between species

Next, we assessed differential gene expression between species within each developmental stage and tissue type (Appendix S6). The lowest number of DE genes between species was in stage 1 distal tissue (6,413 genes) and the highest in stage 4 proximal tissue (7,994 genes; Fig. 6A). Except for stage 2, more genes were DE between species in proximal tissue than distal tissue. All DE genes across stages and tissue types had a logFC between -17.4 and 15.1 (Fig. 6B).

**Fig. 6.**
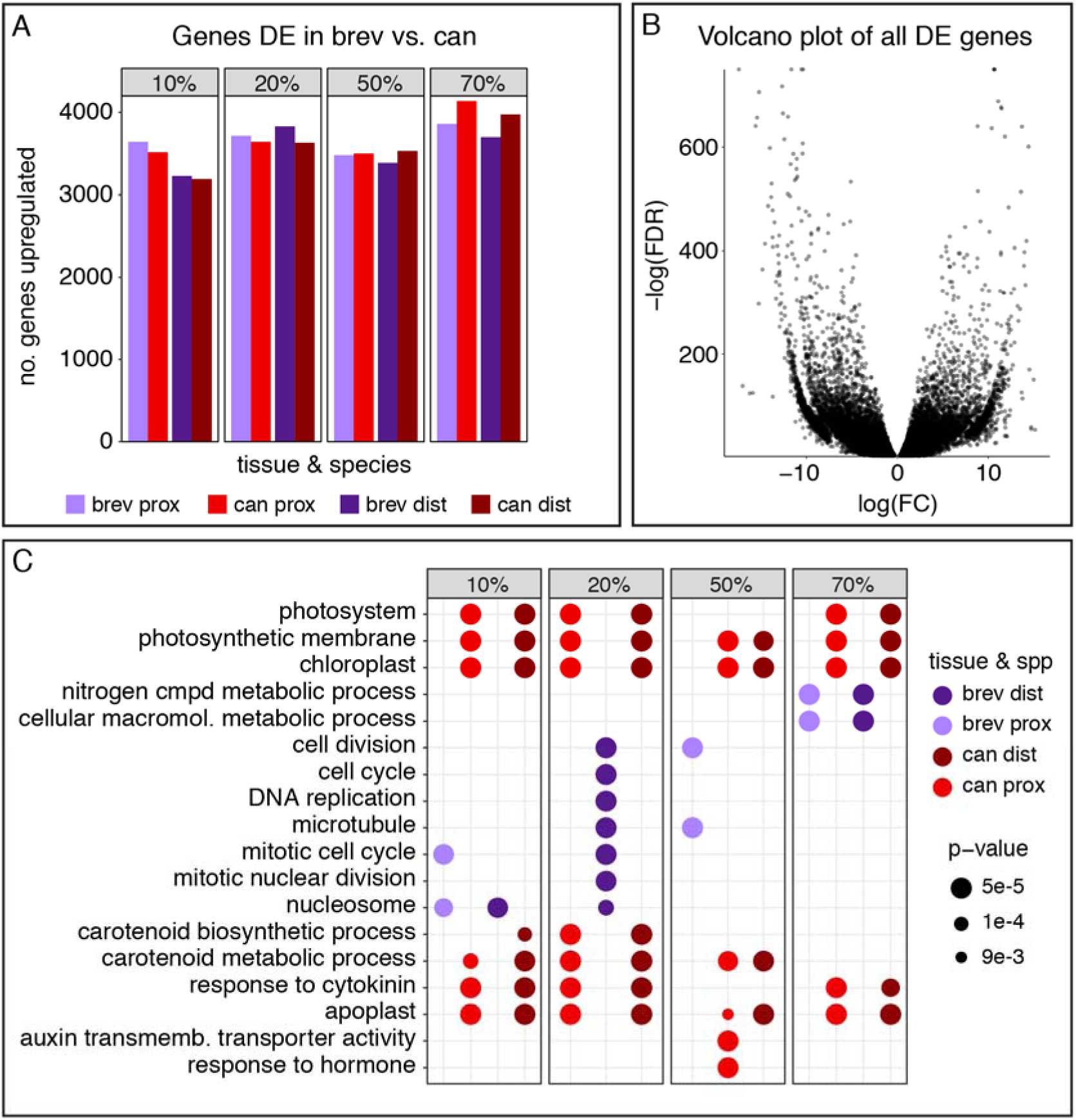
Species DE analysis. **A.** Number of genes DE between *A. brevistyla* and *A. canadensis* within each stage and tissue type. **B.** Volcano plot of all significantly DE genes showing the logFC and -logFDR of each transcript. Points with a positive logFC represent genes upregulated in *A. canadensis*, points with a negative logFC represent genes upregulated in *A. brevistyla*. **C.** Selected GO terms enriched in each species. Abbreviations: brev, *A. brevistyla*; can, *A. canadensis*; prox, proximal; dist, distal.

GO terms for genes upregulated in *A. canadensis* were dominated by photosynthetic processes, although GO enrichment for carotenoid synthesis, hormone signaling, and cell wall development were present as well. Genes upregulated in *A. brevistyla* were enriched for general cellular metabolism and cell division GO terms (Fig. 6C, Appendix S7). Top DE hits varied among all stages and tissue types, but were generally annotated as genes involved in various cellular metabolic functions.

### Differential expression analysis between proximal and distal spur compartments

Finally, we compared expression between the proximal and distal spur compartments within each species and developmental stage. The number of genes significantly DE between proximal and distal compartments ranged from 1,948 in *A. brevistyla* stage 3 spurs to 4,593 in *A. canadensis* stage 2 spurs (Fig. 7A, Appendix S8). More genes were DE in *A. canadensis* than *A. brevistyla,* with the exception of stage 4 spurs, and there were more genes upregulated in the distal compartment than proximal compartment across all stages and species. Across species and stages, the logFC of all significantly DE genes ranged from -4.0-3.5, with several outliers from 4.1-8.9 (Fig. 7B).

**Fig. 7.**
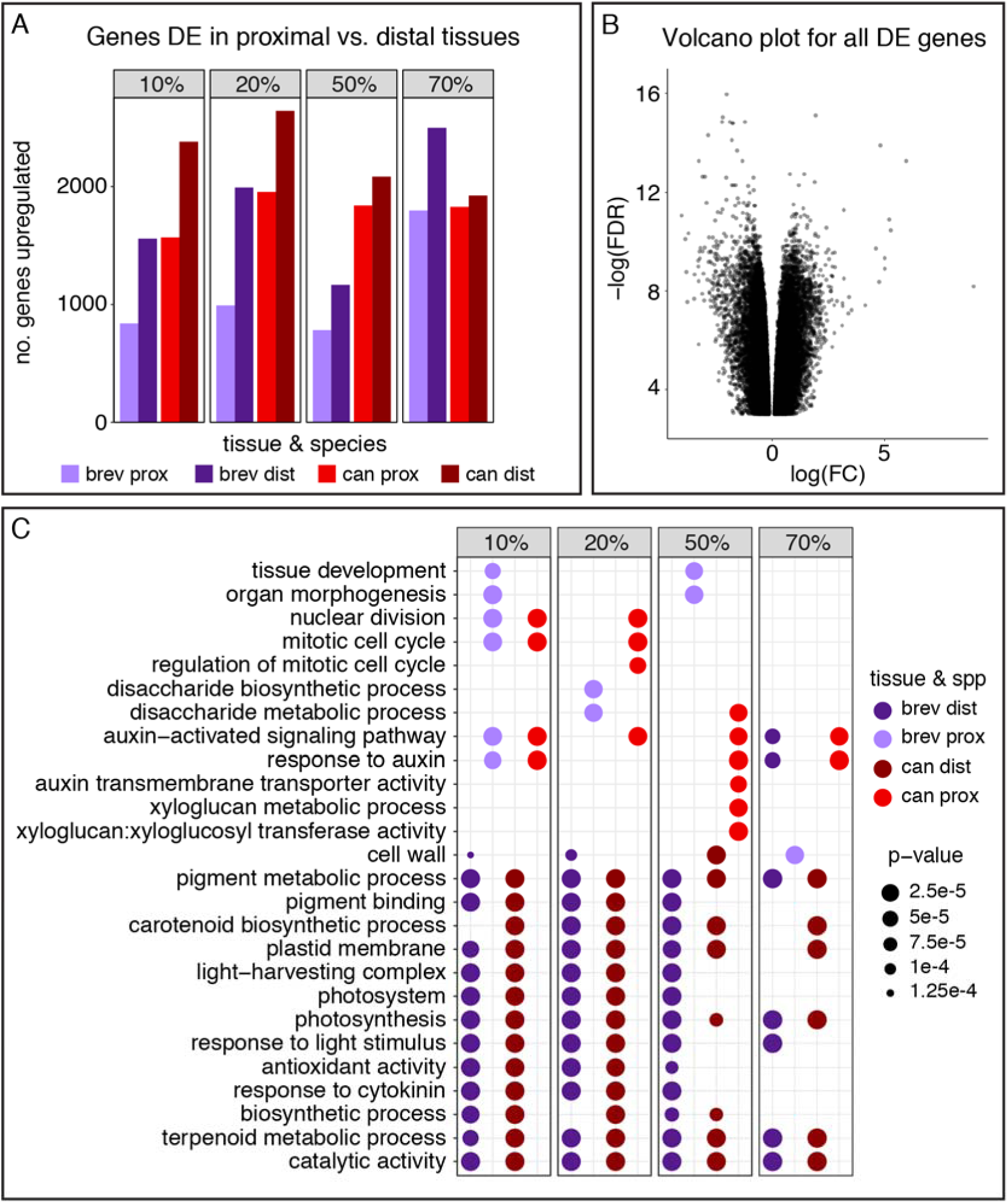
Proximal vs. distal tissue DE analysis. **A**. Number of genes DE between proximal and distal tissues within a species and developmental stage. **B.** Volcano plot of all DE genes showing the logFC and -logFDR of each transcript. Points with a positive logFC indicate genes upregulated in the proximal compartment, and those with a negative logFC indicate genes upregulated in the distal compartment. **C**. Selected GO terms enriched in proximal and distal tissues. Abbreviations: brev, *A. brevistyla*; can, *A. canadensis*; prox, proximal; dist, distal.

Early stage proximal tissue in both *A. canadensis* and *A. brevistyla* was enriched for GO terms related to mitosis and cell division, and mid-stage proximal tissue was enriched for disaccharide metabolic processes (Fig. 7C, Appendix S9). Unique to *A. canadensis* mid- and late-stage proximal tissue was enrichment for auxin signaling and xyloglucan metabolic processes. Distal tissue across stages and species was enriched for GO terms involving photosynthesis, metabolism, and pigment synthesis. Cytokinin response GO terms were enriched in early stage distal tissue.

One of the top hit DE genes among all stages of *A. canadensis* and *A. brevistyla* is the transcript Aqcoe3G370100.1, annotated as a member of the TCP gene family. Protein sequence synapomorphies place it in the CIN-like class II TCPs (Liu et al., 2019), which are known cell division repressors in plant lateral organs (Huang and Irish, 2015; Yant et al., 2015; Danisman, 2016; van Es et al., 2018a). Another gene that stood out as upregulated in later-stage *A. canadensis* proximal tissue, but not DE between proximal and distal tissue in any stage of *A. brevistyla,* was annotated as *INDOLE ACETIC ACID 9* (*IAA9),* which is a member of a gene family that functions downstream of auxin to regulate homeostasis of the hormone response pathway, often in the context of vascular development (Wang et al., 2005; Xu et al., 2019).

### Weighted gene correlation network analysis (WGCNA)

For a more holistic approach to understanding the transcriptional dynamics across our complex sampling scheme, we conducted WGCNA on all expressed genes to identify gene co-expression modules. The analysis yielded 30 modules (MDs), the smallest of which contained 34 genes (MD21) while the largest contained 4,316 genes (MD23; Fig. 8A, Appendix S10). 191 genes of the 21,804 used in the WGCNA were not assigned to any module. To determine potential roles of these modules in spur development, we tested them for correlations with spur developmental stage, proximal/distal tissue type, and species (Fig. 8A) and plotted the mean z-score across all biological replicates within a species, stage, and tissue type for each module (Fig. 9). Many modules showed a clear developmental trend. MDs 20-25 show positive correlations with the earliest developmental stage, the strongest of which is MD23 (0.84, P = 5e-21), and become increasingly negatively correlated with later stages (Fig. 8A). MDs 4-9 generally show the opposite pattern, becoming more positively correlated with later developmental stages, with MD8 showing the strongest correlation with the 70% stage (0.86, P = 7e-23). MDs 26, 27, and 29 exhibit more varied correlation through development, peaking at the 50% stage and then decreasing again. The z-score boxplots generally reflect these trends and reveal more nuanced patterns within species and tissue type (Fig. 9). The analysis also indicated that certain modules were strongly correlated with species, such as MD14 with *A. canadensis* (0.99, P = 5e-61) and MDs 25 and 30 with *A. brevistyla* (0.86, P = 5e-23; 0.82, P = 3e-19, respectively). However, the z-score boxplot of MD14 revealed that although the expression of *A. canadensis* samples is consistently slightly higher than that of *A. brevistyla*, presumably leading to the strong correlation with *A. canadensis,* the two species are quite similar in overall expression pattern within MD14 (Fig. 9). Finally, the distal and proximal traits did not exhibit strong correlations with any modules. The strongest association for the distal compartment was with MD16 (0.25, P = 0.04) and the strongest association for the proximal compartment was with MD7 (0.23, P = 0.05), with both modules showing stronger correlations with developmental stages and species than tissue type (Fig. 8A). However, examination of the z-score boxplots reveals that several modules reflect species-specific differences in compartment expression, such as MDs 5, 7, and 17, which all show high expression in late-stage *A. canadensis* proximal tissue.

**Fig. 8.**
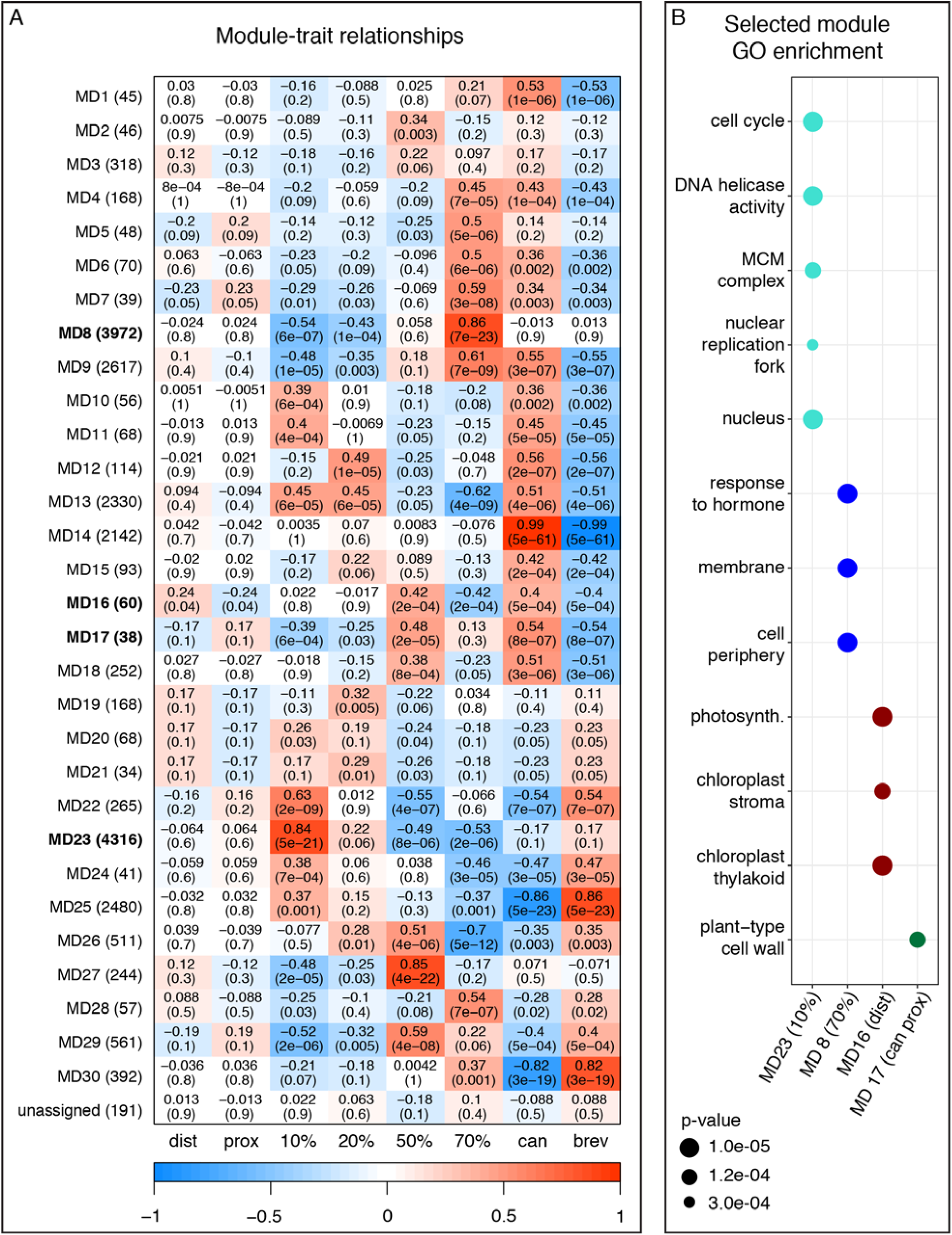
WGCNA results. **A.** Gene co-expression modules identified via WGCNA and their association with sample traits. Modules (MDs) are identified by number on the y-axis, and the number of transcripts in each module is in parentheses. Bolded modules are further explored in B. The sample traits are on the x-axis (dist, distal; prox, proximal; stages 10%-70%; can, *A. canadensis*; brev, *A. brevistyla*). The correlation between module and sample trait is indicated by the heat map; the top number in each heat map cell represents the strength of the correlation, and the bottom number in parentheses is the p-value. Red cells indicate a strong positive correlation, and blue cells indicate a strong negative correlation. **B.** Selected GO-term enrichment for MD23 (10% stage-associated), MD8 (70%), MD16 (dist), and MD17 (can prox) modules.

**Fig. 9.**
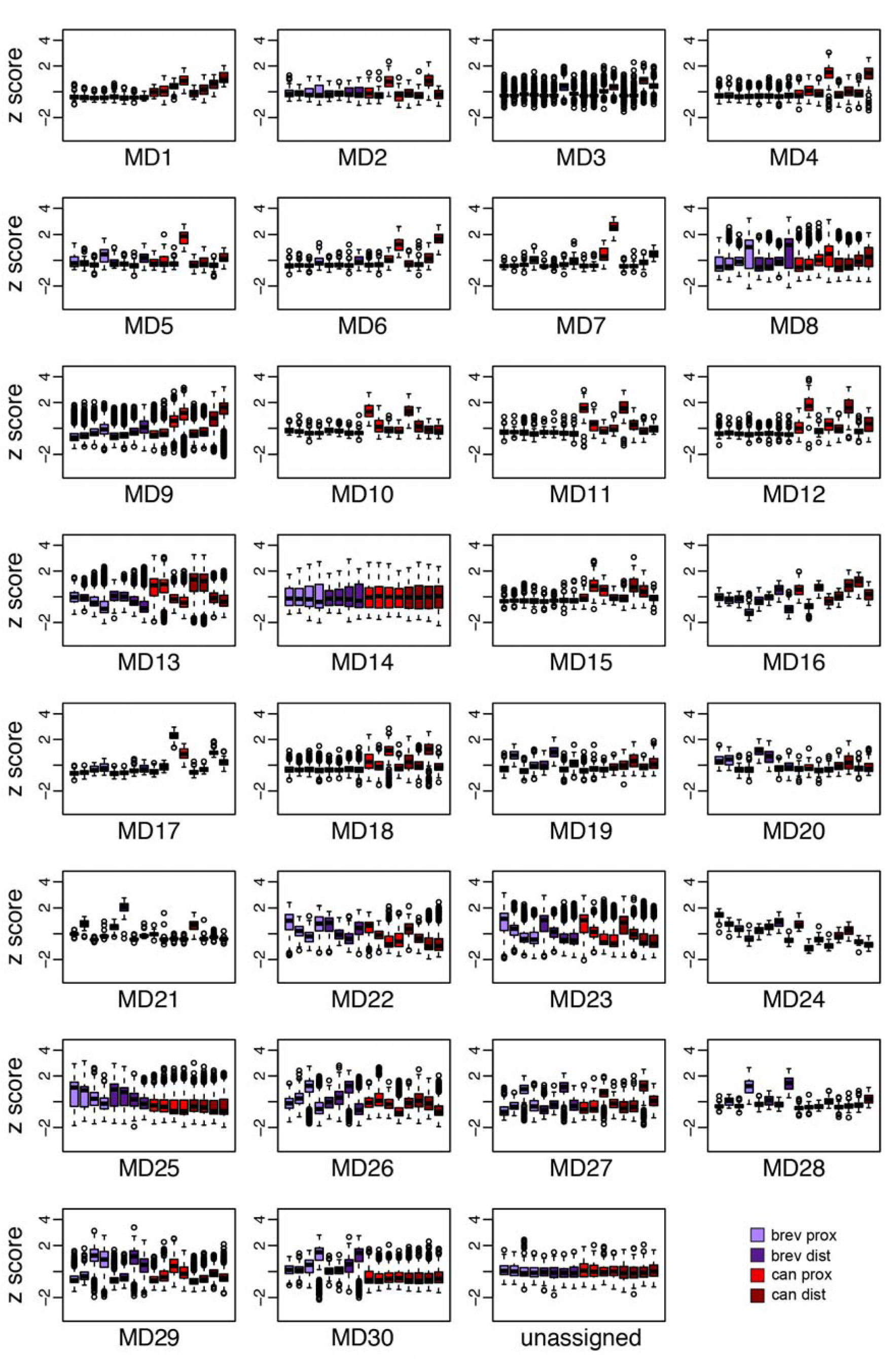
Box plots of the mean z-score of all biological replicates per species, developmental stage, and tissue type genes in each WGCNA module (MD) as well as those unassigned to any module. *A. brevistyla* proximal tissue (brev prox) stages 10%-70% are represented in light purple (first four samples) and distal tissue (brev dist) stages 10%-70% in dark purple (second four samples); *A. canadensis* proximal tissue (can prox) stages 10%-70% in light red (third four samples), and distal tissue (can dist) stages 10%-70% in dark red (last four samples).

We performed GO-term analyses on modules showing strong correlations with single traits or tissue-specific expression patterns as revealed by the z-score plots and compared module genes to the DE genes from our previous analyses (Fig 8B, Appendix S11). MD23 (10% stage-associated) was enriched for terms related to cell division, and 49.4% of the genes in the module are also upregulated in the 10% stage in the early vs. late DE analysis. MD8 (70% stage) and MD27 (50% stage) were enriched for cell wall, cell membrane, and hormone signaling GO terms, and 47.6% of the MD8 genes are upregulated in the 70% stage in the early vs. late DE analysis. The species-associated modules exhibited next to no GO enrichment. *A. brevistyla*- associated MD30 did not have any significant GO enrichment, and MD25 (also *A. brevistyla*- associated) only returned a single significant GO term for ADP binding. However, 88.7% of the MD30 genes and 66.3% of the MD25 genes are also upregulated in *A. brevistyla* in the species DE analysis. MD14 (*A. canadensis-*associated) was also only enriched for the ADP binding GO term, and 44.2% of its genes are upregulated in *A. canadensis*. The weakly distal-associated MD16 was enriched for GO terms related to photosynthesis and had an 81.7% overlap with genes upregulated in the distal compartment in the proximal vs. distal DE analysis. While the canadensis proximal-associated MDs 5 and 7 did not exhibit significant GO enrichment, MD17 had a single significant GO term for plant-type cell wall (Fig. 8B) and a 68.4% overlap with genes upregulated in *A. canadensis* proximal tissue. MD17 contains multiple genes related to cell wall synthesis, carbohydrate metabolism, and xylem development.

### Exploration of candidate genes of interest

We recently identified QTL contributing to spur length and curvature variation between *A. canadensis* and *A. brevistyla* (Edwards et al., 2021). Since the developmental data from the present study revealed that cell number is responsible for the spur length differences between the species, we looked for genes involved with cell division within the six spur length QTL that we previously identified. The 1.5 LOD interval of the QTL with the largest effect on spur length on chromosome 7, responsible for 16.1% of the trait variance, contains 457 genes, 350 of which are expressed in the transcriptomic dataset (cpm ≥1 in at least 5 samples; Appendix S12). These include transcripts annotated as *GROWTH REGULATING FACTOR 5 (GRF5),* a cell division promoter that is associated with petal shape and size variation in *Chrysanthemum* cultivars (Ding et al., 2019); *EUKARYOTIC TRANSLATION INITIATION FACTOR 4A1 (EIF4A1),* which affects cell cycle progression via protein translation regulation (Bush et al., 2015, 2016); *CYCLOPHILIN71 (CYP71)*, a WD40 domain cyclophilin that targets genes related to lateral organ development and meristem activity in *Arabidopsis* (Li et al., 2007); and *AURORA1 (AUR1),* a kinase associated with cell division machinery that peaks in activity during mitosis (Demidov et al., 2009). While *CYP71* is expressed at low levels, the others are expressed at moderate to high levels that decrease in later developmental stages (Appendix S13). The 1.5 LOD interval of the single locus of large effect for spur curvature (responsible for 46.5% of spur curvature variance) is also on chromosome 7, but does not overlap with the interval for spur length. It contains 199 genes (Appendix S14), of which 144 are expressed in the RNA-seq dataset and 117 are DE between species and/or between proximal and distal spur compartments for at least one developmental stage. The developmental basis for spur curvature is more complex than that of spur length, but we would expect relevant loci to play some role in the regulation of cell elongation, either promoting it in *A. canadensis* or repressing it in *A. brevistyla.* Few of the 144 expressed genes in the 1.5 LOD interval have clear roles in cell expansion, but there is one locus, Aqcoe7G339000, that encodes *IBL1,* a bHLH transcription factor known to negatively regulate cell elongation (Fig. 10A; Zhiponova et al., 2014).

**Fig. 10.**
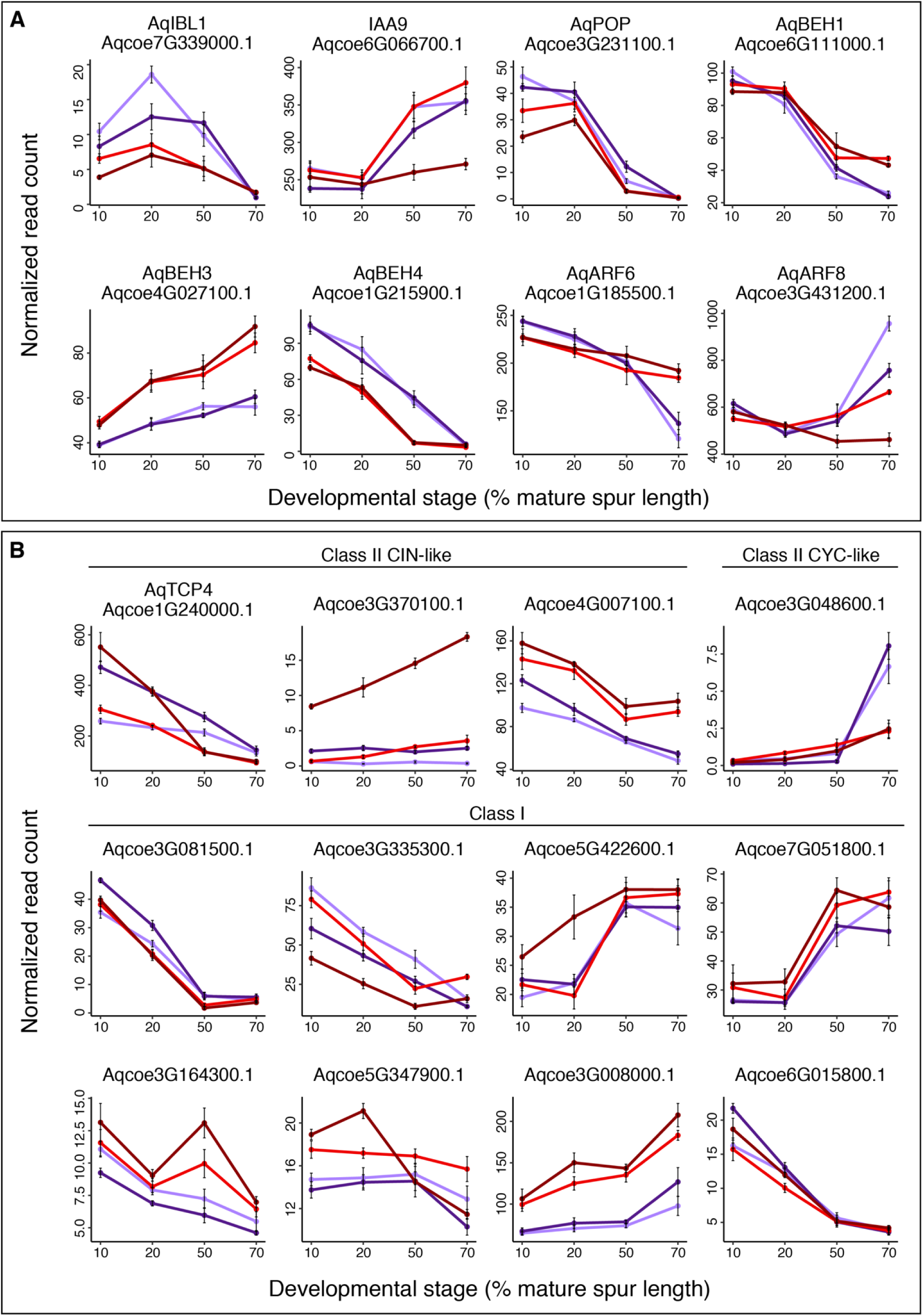
Expression patterns of genes of interest by species, tissue, and developmental stage. Mean normalized read counts across all biological replicates are shown, with error bars representing the standard error of the mean. Light purple, *A. brevistyla* proximal tissue; dark purple, *A. brevistyla* distal tissue; light red, *A. canadensis* proximal tissue; dark red, *A. canadensis* distal tissue. **A.** Expression patterns of genes of interest identified by the present study as well as previous studies in *Aquilegia*. **B.** Expression patterns of all TCP family genes in the transcriptomic dataset, organized by TCP subfamily.

While our QTL for spur length and curvature harbor the major genetic variants influencing divergence in these traits, many additional loci with smaller effects likely also contribute to variation in these traits. To explore additional candidates outside of our major QTL, we plotted the average normalized expression of several genes of interest to see if they have tissue-, species-, or stage- specific expression patterns that would indicate involvement in controlling spur morphology in our system (Fig. 10A). *IAA9* emerged from the present study, while the others are known regulators of *Aquilegia* nectar spur cell division (*AqPOP*; Ballerini et al., 2020) or cell elongation (*AqBEH*s and *AqARF*s; Zhang et al., 2020; Conway et al., 2021). Although several of these loci show differential expression between species, only *AqARF8* shows dramatically different expression between the distal and proximal compartments in both species, while *IAA9* shows high differential expression between compartments only in *A. canadensis* (Fig. 10A).

Given that a TCP-family transcription factor was consistently a top DE hit across our proximal vs. distal analyses, and that *AqTCP4* is a known regulator of cell proliferation in the *Aquilegia* spur with a distal-specific phenotype in *A. coerulea* ‘Origami’ (Yant et al., 2015), we particularly wanted to explore the expression of the TCPs more broadly. We identified all expressed genes in our dataset annotated with the TCP protein family ID PF03634, assigned them to one of the three TCP subfamilies (Class I, Class II CIN-like, or Class II CYC-like) based on known sequence synapomorphies (Liu et al., 2019), and plotted their expression patterns (Fig. 10B). These plots confirm the general distal bias of Class II CIN-like homologs, but also highlight an interesting Class I homolog, Aqcoe3G335300.1, which appears to have a proximal bias.

## DISCUSSION

### Development & evolution of contrasting spur morphologies

Organ curvature requires differential growth between the inner and outer surfaces of the curve. Before determining the developmental mechanisms underlying *Aquilegia* spur curvature, we first compared the extent of differential growth in spur surfaces across species with contrasting petal morphologies. Unsurprisingly, we confirmed that curved-spurred *A. brevistyla* and *A. saximontana* exhibit significantly different proximal and distal spur surface lengths, while straight-spurred *A. formosa* and *A. canadensis* exhibit little to no differential growth between spur surfaces. Next, we sought to understand the nature of this differential growth in curved spurs. In mature petals of both *A. brevistyla* and *A. saximontana,* we found a consistent pattern of increased cell numbers in the distal spur surface, suggesting that this could be a conserved pattern for the ancestral bee-pollinated, curved spur. Our analysis of multiple developmental stages of *A. brevistyla* spurs revealed that a longer duration of cell division in the distal surface is responsible for its greater cell number. These findings align with what we observe in the gross morphology of early flower buds: spurs are visibly curved extremely early in development, when cell divisions are active (Yant et al., 2015; Ballerini et al., 2019). If spur curvature were caused solely by differential cell elongation, we would expect it to appear later in development when cell elongation is the primary driver of growth (Puzey et al., 2012). We then turned to the question of how straight spurs evolved during the transition to hummingbird pollination. We were surprised to find that instead of equalizing cell number between proximal and distal surfaces, differential cell divisions were maintained in *A. canadensis,* and straightening occurred via differential cell elongation. The fact that we found the same pattern in *A. formosa,* which convergently evolved hummingbird pollination, suggests there may be aspects of the spur developmental program that constrain this evolutionary process. These results are consistent with the *AqTCP4*-silencing phenotype of *A. coerulea* ‘Origami’, which revealed that the proximal and distal compartments of these straight spurs are still functionally dissociable (Yant et al., 2015). It appears that transitioning from curved to straight spurs does not occur by eliminating the differential control of these two domains and, in fact, involves employing distinct developmental mechanisms in each compartment.

Although we observed differences in cell number between the proximal and distal spur surfaces in both *A. brevistyla* and *A. canadenis* spurs at maturity, when we examined earlier developmental stages, we found that the difference is established in a contrasting manner between the two species*. A. canadensis* spur surfaces appear to be experiencing differential *rates* of cell division, in contrast to the differential *duration* of cell division that occurs in *A. brevistyla*. By the earliest stage we could sample in *A. canadensis,* there is already a significant difference in cell number between the proximal and distal surfaces, suggesting that the mitotic rate difference between surfaces in this species is established early in development, while a significant difference in cell number between surfaces is not apparent in *A. brevistyla* until the 30% stage. These results suggest that although differential cell division between surfaces appears to have been “conserved” during the transition from bee- to hummingbird-pollination, the exact developmental mechanism by which it is achieved may have changed. Fascinatingly, the 10% stage spur in *A. canadensis* contains twice the number of cells as the 20% stage spur in *A. brevistyla,* even though their total spur lengths are roughly the same at these stages (Fig. 3A, Appendix S1). This reveals how early the overall spur cell number difference is established between species and raises fundamental questions about how developmental processes occurring in the nascent spur affect final spur morphology – does *A. canadensis* initially recruit more cells to its spur than *A. brevistyla*, or simply experience dramatically faster rates of cell division? These species differences are potentially further complicated when considering overall organ size. *A. brevistyla* is commonly known as the small-flowered columbine and its spurs are significantly smaller than the other taxa examined here, especially *A. canadensis* (Appendix S1).

Spur length varies dramatically throughout the genus, ranging from 1cm in bee-pollinated species like *A. brevistyla* to 16cm in *A. longissima* (Munz, 1946). Previous work that explored a portion of this spur length diversity found that it was closely correlated with the Phase II process of anisotropic cell elongation (Puzey et al., 2012). However, the current study uncovered clear variation in cell number. *A. brevistyla,* a species with one of the shortest spurs in *Aquilegia* (Munz, 1946), has approximately half the number of spur cells compared to the other three species, which all have spurs roughly twice as long as *A. brevistyla’s.* Cell length, on the other hand, is comparable across all four species, and *A. brevistyla* even has the longest distal surface cells of the group. These differences with Puzey *et al*. are most likely due to species sampling. Our study focused on the transition from bee- to hummingbird-pollination and included spurs at the shorter extreme of the length distribution (e.g., *A. brevistyla*). In contrast, for the developmental cell-counting analysis, Puzey *et al*. examined relatively longer-spurred taxa – *A. canadensis* (∼2.5cm, hummingbird-pollinated)*, A. coerulea* (∼6.5cm, hawkmoth), and *A. longissima* (16cm, hawkmoth). Taken together, these datasets paint a complex picture of spur morphological diversification in *Aquilegia.* It appears that multiple developmental mechanisms have contributed to the full breadth of spur lengths in the genus, with variation in cell number associated with the shorter extremes, and variation in anisotropic expansion associated with longer extremes. Likewise, both cell number and length contribute to the generation of curvature and the evolution of straightening.

A key next step will be extrapolating our largely two-dimensional datasets to a three-dimensional model, which should be possible using recently developed live-imaging approaches (Min et al., 2021) combined with increased sampling across the genus. These developmental findings also shed new light on our spur morphology genetic architecture data for *A. brevistyla* and *A. canadensis* (Edwards et al., 2021). Promising candidate genes for spur length differences between these taxa will likely be involved in cell proliferation, as overall cell number seems to drive spur length in this comparison, while candidates for spur curvature could control cell elongation, as differential cell elongation is the primary developmental difference between straight and curved spurs.

### Synthesis of developmental & transcriptomic data

At multiple levels of comparison – developmental stage, species, and tissue type – the transcriptomic data consistently validate observations from the spur development analyses and provide key insights into the genetic dynamics underlying the transition from curved to straight spurs. These comparisons also yielded similar metabolic noise, such as the prevalence of photosynthesis GO term enrichment in early, distal, and *A. canadensis* tissues; petals are still green at the 10% developmental stage, and distal and *A. canadensis* tissues have increased light exposure compared with inward-facing proximal tissue and shorter *A. brevistyla spurs*. Noise aside, there is clear and meaningful developmental signal to be explored in this multifaceted transcriptomic dataset.

In general, the comparison of the earliest (10%) and latest (70%) developmental stages was entirely consistent with previous studies (Ballerini *et al*., 2019; Puzey *et al*. 2012; Yant *et al*. 2015) in detecting early cell division and late cell expansion. The large number and logFC magnitude of DE genes that came out of this analysis reflect the dramatic changes occurring in the spur between these stages. The substantial overlap in gene lists between the DE analyses and stage-associated WGCNA modules, as well as their similarity in GO enrichment, indicates that there is a strong and consistent evidence of developmental maturation across species and tissue types.

Comparisons of gene expression between *A. canadensis* and *A. brevistyla* also yielded many highly differentially expressed genes, and while the insight they provide into the transcriptional basis of spur length differences between the species is somewhat limited, they reveal a potentially key molecular process underlying differences in spur shape. As shown by our developmental study, the overall cell number difference between *A. canadensis* and *A. brevistyla* spurs is already established by the first stage of our sampling scheme, therefore its transcriptional signal may not have been directly captured by our dataset. However, it is notable that the first three stages of *A. canadensis* tissue are enriched for GO terms associated with the cytokinin response, a known promoter of cell division (Hwang et al., 2012; Li et al., 2021). The transcriptomic data also validate another observation related to cell proliferation from our developmental data: genes upregulated in *A. brevistyla* are consistently enriched for cell division GO terms, not because *A. brevistyla* spurs have a higher number of cells than *A. canadensis*, but because cell divisions are continuing in *A. brevistyla* long after they have leveled off in *A. canadensis*.

The transcriptional signal of spur shape between species was clearer than that of spur length. The main developmental difference between *A. brevistyla* and *A. canadensis* that appears to be responsible for the straightening of the spur is increased differential cell elongation between spur compartments in *A. canadensis.* Genes upregulated in *A. canadensis* 50% stage proximal tissue (when and where increased rate of cell elongation becomes prominent) are enriched for auxin signaling GO terms, a process associated with cell expansion in above-ground plant organs (Went, 1927; dela Fuente and Leopold, 1970; Christian et al., 2006) and specifically in *Aquilegia* petal spurs (Zhang et al., 2020).

Of the three levels of comparisons we examined, the proximal vs. distal analysis yielded the smallest and lowest-magnitude DE gene lists, and the overall proximal/distal trait associations with the WGCNA modules were much weaker than with species or developmental stage traits. However, given that we were comparing tissue from within the same organ, the number of DE genes was surprisingly high, further validating preliminary evidence that the proximal and distal compartments are experiencing very different developmental programs. Moreover, closer examination of the z-score boxplots of the WGCNA modules revealed that many modules had tissue-specific expression patterns within developmental stages and/or species. As a whole, the proximal-distal comparison provided valuable insight into the genetic underpinnings of contrasting *Aquilegia* spur morphologies.

Our developmental study showed that the spurs of both species experience greater cell division in their distal surfaces than proximal, and the cytokinin response GO term is enriched in *A. canadensis* distal tissue through the 20% stage, and in *A. brevistyla* through the 50% stage, corresponding to the cessation of distal cell division in each species. Given cytokinin’s role in promoting cell proliferation (Hwang et al., 2012; Li et al., 2021), the elevated cytokinin response in the distal surface could be the transcriptional signal of the differential cell division observed in the developmental data. However, there are also many GO terms related to mitosis and the cell cycle enriched in early-stage proximal tissues in both species. While this seems to contradict the developmental data at first glance, we believe it can be explained by the fact that the nectary, which undergoes prolonged cell division to produce its tightly-packed secretory cells (Antoń and Kamińska, 2015; Yant et al., 2015), is positioned towards the proximal spur surface, leading to an uneven distribution between the proximal and distal dissections. The expression profiles of *STYLISH1 (AqSTY1)* and *AqSTY2*, which specifically control nectary development in *Aquilegia* (Min et al., 2019), support this theory, showing upregulation in early-stage proximal tissues in both species (Appendix S15). Disaccharide biosynthesis and metabolism GO terms, which are relevant to nectar production, are also enriched in proximal tissue in this dataset.

These analyses further solidify our understanding of the transcriptional underpinnings of differential cell elongation in the spur in relation to spur shape. Genes upregulated in late-stage *A. canadensis* proximal tissue compared to distal tissue are enriched for auxin-related GO terms, while those upregulated in *A. brevistyla* proximal tissue compared to distal are not, echoing the results from the species-level comparisons. The fact that both the proximal-distal and between-species analyses highlight auxin signaling in the *A. canadensis* proximal spur compartment is robust evidence in support of the auxin’s possible role in the differential cell elongation process that is critical to developing straight spurs.

### Promising candidate genes for future study

We used our discoveries about the developmental and transcriptional underpinnings of spur morphological differences in *A. canadensis* and *A. brevistyla* to inform our search for promising candidate genes for future evaluation. We first turned to the QTL mapping data for spur length and curvature from Edwards *et al*. (2021). The genetic architecture of spur length is the more complex of the two traits, consisting of 6 loci of small- to moderate-effect. Our developmental study revealed that differential cell proliferation between *A. canadensis* and *A. brevistyla* is the mechanism underlying this trait difference, enabling us to focus our search for potential causative genes on known players in regulation of cell division. Interestingly, *AqPOP,* a known mitotic regulator in the spur, is not in the 1.5 LOD interval of the minor locus on chromosome 3. However, we have identified several genes under the largest spur length QTL on chromosome 7 that bear further investigation. Although our transcriptional sampling does not cover the extremely early stage during which cell number differences between species are established, we still believe they are worthy of further investigation due to their known roles in cell division regulation in other systems (Li et al., 2007; Demidov et al., 2009; Bush et al., 2015, 2016; Ding et al., 2019).

Spur curvature, on the other hand, has a simpler genetic architecture consisting of only a single locus of large effect on chromosome 7, and from our developmental study we knew to look for known regulators of cell elongation in the 1.5 LOD interval of this locus. While almost 200 genes from the transcriptional study were expressed in this interval, and a significant fraction of these were DE between species and/or tissues, a bHLH transcription factor stands out as a possible candidate. Aqcoe7G339000 encodes a transcript annotated as a homolog of *IBL1 (INCREASED LEAF INCLINATION BINDING BHLH1- LIKE),* so named for its similarity to *INCREASED LEAF INCLINATION BINDING BHLH1 (IBH1)* (Zhiponova et al., 2014). In rice, INCREASED LEAF INCLINATION BINDING 1 (ILI1) and IBH1 antagonize each other in regulating cell elongation downstream of the brassinosteroid (BR) pathway, as do their respective homologs in Arabidopsis, PACLOBUTRAZOL RESISTANCE 1 (PRE1) and AtIBH1 (Zhang et al., 2009). ILI1/PRE1 promote cell elongation while IBH1 and its paralog IBL1 repress it, integrating signals from the BR and auxin pathways (Zhang et al., 2009; Ikeda et al., 2012; Bai et al., 2013; Zhiponova et al., 2014). In our transcriptomic data, the *AqIBL1* locus is significantly upregulated in *A. brevistyla* proximal tissue compared to distal tissue in early stages (Fig. 10A), but in *A. canadensis* is expressed at much lower levels than *A. brevistyla* and shows little to no DE between tissue types. Previous studies in *Aquilegia* have confirmed a key role for both the BR and auxin pathways in promoting spur cell elongation (Zhang et al., 2020; Conway et al., 2021). Taken together, this suggests a model in which expression of *AqIBL1* could repress cell elongation in the proximal compartment of the curved *A. brevistyla* spur, while reduction of expression in *A. canadensis* relieves the repression, allowing BR and auxin to promote cell elongation and thereby straighten the spur. This model is also consistent with the implication of auxin signaling as playing a role in *A. canadensis* cell elongation since *AqIBL1* would be expected to function in this pathway. Functional characterization of this locus is a key goal, but conducting such studies in curved spur species is challenging, in part because we only have transient gene knock-down techniques in *Aquilegia.* Additionally, the chromosome 7 QTL only explains 46.5% of the F2 variance, which may yield relatively weak phenotypes. However, further exploration of the ILI/ILH regulatory program as well as the contributions of auxin and BR signaling to spur curvature are certain to provide important insights.

Of course, there is still 53.5% of spur curvature that appears to be explained by QTL too small to be detected in the 2021 analysis. Our new developmental and transcriptional data combined with previous work in *Aquilegia* have led us to additional candidates that could be contributing to the other 53.5% of spur curvature. A promising candidate is the transcript annotated as *IAA9,* which was consistently upregulated in *A. canadensis* proximal tissue in later stages, but not DE between the two tissue types at any stage in *A. brevistyla* (Fig. 10A). *IAA9* function has been explored both in *Populus tomentosa* and *Solanum Lycopersicon*, where it plays diverse roles in programs regulated by auxin, including vascular development, leaf architecture, and fruit set (Wang et al., 2005; Xu et al., 2019). This finding is consistent with a potential role for auxin in spur development, particularly cell elongation, but also illuminates a possibility that we have not yet explored, the role of vasculature in the evolution of spur morphology. A key next step in our study of spur development will be comprehensive imaging of the developing spur that can go beneath the epidermal tissue layer, allowing us to determine whether and how differences in vascular architecture affect the gross morphology of the nectar spur.

There have also been numerous previous characterizations of *Aquilegia* spur cell proliferation and elongation regulators in straight-spurred *A. coerulea* ‘Origami.’ While *AqPOP,* the *AqARFs*, and the *AqBEHs* do not immediately stand out as potential candidates for influencing spur curvature based on their expression patterns in *A. canadenis* and *A. brevistyla*, the *AqARFs* may be worth investigating based on the transcriptomic evidence that auxin signaling could be playing a role in spur curvature. Additionally, the ARFs and BEHs are post-transcriptionally regulated, so their potential role cannot be assessed on expression pattern alone. *AqTCP4,* however, was the first gene to indicate how genetically distinct the proximal and distal spur compartments might be, and the TCP family genes appeared as DE across our transcriptomic analyses. We confirmed the distal bias of *AqTCP4,* and discovered that this is a pattern shared among the Class II CIN-like TCPs, which are known regulators of floral development and often repress cell division and enhance cell differentiation (Fig. 10B; Efroni et al., 2008; Huang and Irish, 2015; van Es et al., 2018). Although class II CYC-like TCPs have been shown to regulate petal development and symmetry in *Antirrhinum* (Luo et al., 1996; Corley et al., 2005), the only homolog in our dataset is most likely not an important regulator of spur shape due to its extremely low expression levels. Several Class I TCPs bear further investigation due to their differential expression between species and/or spur compartments, especially Aqcoe3G335300 based on its proximal bias. Class I TCPs play diverse roles in flowering time (Balsemão-Pires et al., 2013), hormone signaling (Danisman et al., 2012; Wang et al., 2015), circadian rhythms (Wu et al., 2016), and development (Weir et al., 2004), so we will not speculate on potential function of specific transcripts here. Further work is needed to collectively assess the role of this important gene family in *Aquilegia* spur development and diversification. It will be especially interesting to compare TCP gene functions in *A. canadensis* and *A. brevistyla* to understand whether and how they contribute to sculpting different spur morphologies.

## CONCLUSIONS

The powerful combination of developmental and transcriptomic analyses of *A. brevistyla* and *A. canadensis* spurs has yielded a wealth of insight into the crucial shift from bee- to hummingbird-pollination in *Aquilegia*, and the rapid radiation of the genus more broadly. Perhaps most surprising is the degree to which development is independently controlled in the proximal and distal compartments of the spur, and the complexity of the developmental processes underlying the diversification of the *Aquilegia* petal. These developmental differences between proximal and distal spur compartments are holistically reflected in our transcriptomic analyses, which enabled us to discern specific molecular processes and candidate genes for further investigation. The potential contributions of auxin signaling, TCP family genes, and vascular architecture to shaping *Aquilegia’s* spur diversity, as well as the identification of possible causative loci from our spur length and curvature QTL maps, are the next stops on an exciting road map for future work in this system. Overall, these studies of *A. canadensis* and *A. brevistyla* have underscored the considerable genetic and developmental intricacy involved in the evolution of this relatively recent pollinator shift, and made the rapid diversification of the genus that much more impressive.

## ACKNOWLEDGEMENTS

The authors thank the Bauer Core Facility at Harvard University for their RNAseq expertise, Adam Freedman at the Harvard FAS Informatics Group for his guidance with data analysis, Stephanie J. Conway for tissue dissection training and support with analysis, Mark Faherty and Mass Audubon for facilitating our *A. canadensis* sampling, and Robin Hopkins, Scott A. Hodges, and Ned Friedman for their thoughtful feedback on the manuscript. This work was supported by a National Science Foundation award IOS-1456217 to E.M.K., and the Department of Organismic and Evolutionary Biology at Harvard University Student Research Award to M. B. E.

## AUTHOR CONTRIBUTIONS

E.M.K. oversaw the study. All experimentation and data collection were conducted by M.B.E. Transcriptomic analysis was performed by M.B.E. with training and oversight by E.S.B. M.B.E. wrote the manuscript with input from the co-authors.

## DATA AVAILABILITY

Raw RNA-seq data are deposited in the Sequence Read Archive under BioProject ID PRJNA818338

## SUPPORTING INFORMATION

Additional supporting information may be found in the following files:

Appendix S1. Petal imaging and cell measurements.

Appendix S2. RNA-seq sampling scheme (continued from Fig. 4A).

Appendix S3. RNA-seq read counts.

Appendix S4. Early- vs. late-stage DE analysis results.

Appendix S5. GO term analysis results for genes DE in early and late stages.

Appendix S6. *A. canadensis* vs. *A. brevistyla* DE analysis results.

Appendix S7. GO term analysis results for genes DE in *A. canadensis* and *A. brevistyla*.

Appendix S8. Proximal vs. distal DE analysis results.

Appendix S9. GO term analysis results for genes DE in proximal and distal tissues.

Appendix S10. Gene membership in WGCNA modules.

Appendix S11. GO term analysis results for WGCNA modules of interest.

Appendix S12. Genes contained in the 1.5 LOD interval of the largest QTL for spur length on chromosome 7.

Appendix S13. Expression patterns of cell division regulatory genes in the 1.5 LOD interval of the chromosome 7 spur length QTL.

Appendix S14. Genes contained in the 1.5 LOD interval of the major QTL for spur curvature on chromosome 7.

Appendix S15. Expression patterns of *AqSTY1* and *AqSTY2*.

