## Appendix S1 for "Complex developmental and transcriptional dynamics underlie pollinator-driven evolutionary transitions in nectar spur morphology in *Aquilegia* (columbine)"

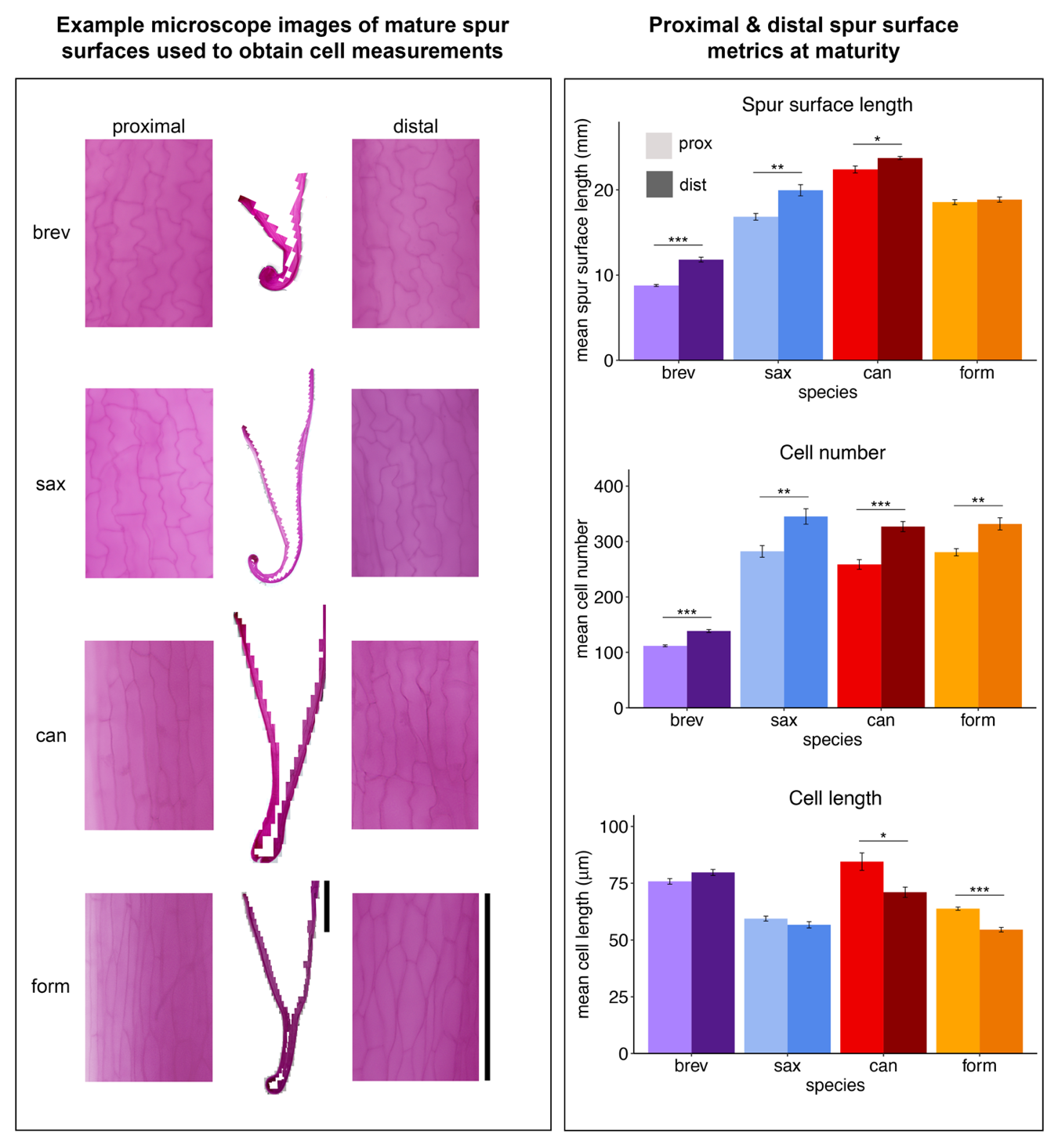

**Appendix S1.** Petal imaging and measurement. **Left:** Imaging. Petals (n=6 per species) were fixed and stained with Acid Fuchsin, then folded longitudinally and flattened between a microscope slide and coverslip for imaging. Brightfield microscope images at 20x were taken of the entire proximal and distal spur surfaces and used to obtain cell count and measurement data. Representative spur panoramas from each species are shown (center), with 130 x 200μm insets of proximal and distal surface cells. Panorama scale bar = 5mm, magnified detail scale bar = 200μm. **Right:** Mean spur metrics across species and proximal/distal compartments. Error bars represent standard error of mean measurements across n=6 spurs per species. P-values of t-test to determine significance of difference between proximal and distal side measurements within a species are denoted with asterisks: *<0.05, **<0.01, ***<0.001 Abbreviations: brev, *A. brevistyla*; sax, *A. saximontana*; can, *A. canadensis*; form, *A. formosa*; prox, proximal; dist, distal.
