## Appendix S2 for "Complex developmental and transcriptional dynamics underlie pollinator-driven evolutionary transitions in nectar spur morphology in *Aquilegia* (columbine)"

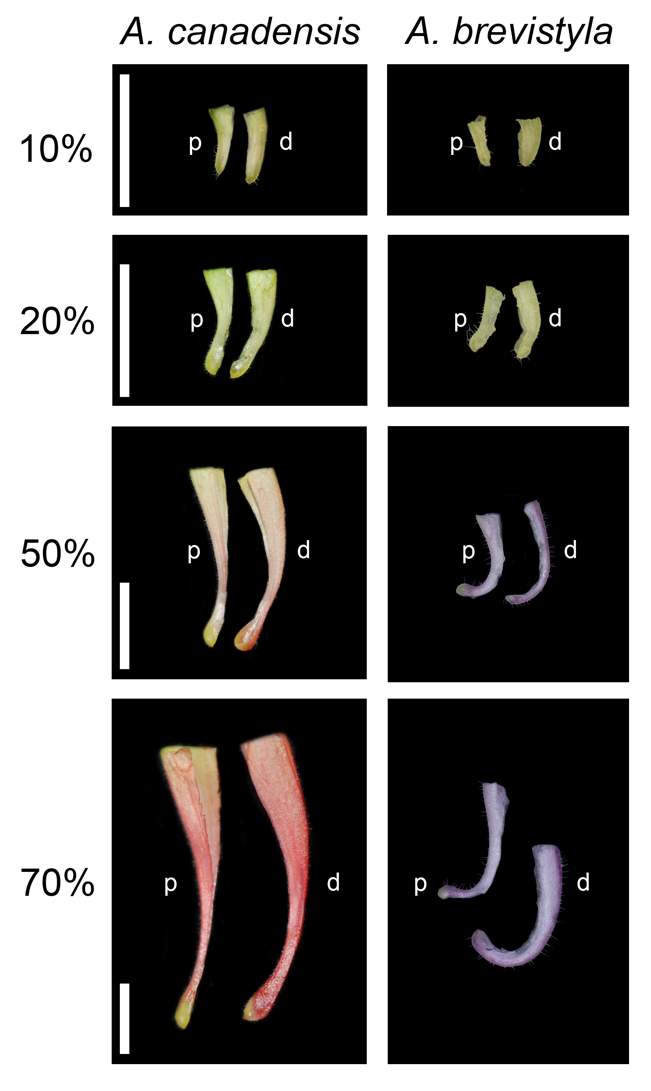

**Appendix S2.** RNA-seq sampling scheme (continued from **Fig. 4A**) showing *A. canadensis* and *A. brevistyla* spurs at four developmental stages dissected into proximal (p) and distal (d) halves. Scale bar = 500μm.
