## Appendix S13 for "Complex developmental and transcriptional dynamics underlie pollinator-driven evolutionary transitions in nectar spur morphology in *Aquilegia* (columbine)"

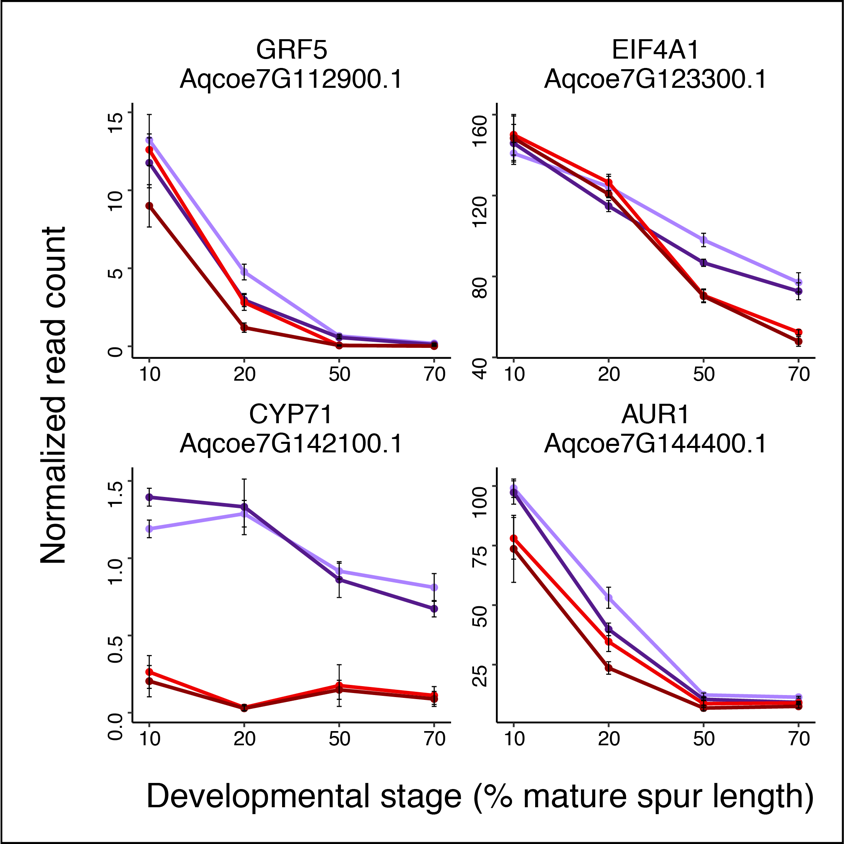

**Appendix S13.** Expression patterns of genes involved in regulation of cell division in the largest QTL for spur length on chromosome 7, by species, tissue, and developmental stage. Mean normalized read counts across all biological replicates are shown, with error bars representing the standard error of the mean. Light purple, *A. brevistyla* proximal tissue; dark purple, *A. brevistyla* distal tissue; light red, *A. canadensis* proximal tissue; dark red, *A. canadensis* distal tissue.
