## Appendix S15 for "Complex developmental and transcriptional dynamics underlie pollinator-driven evolutionary transitions in nectar spur morphology in *Aquilegia* (columbine)"

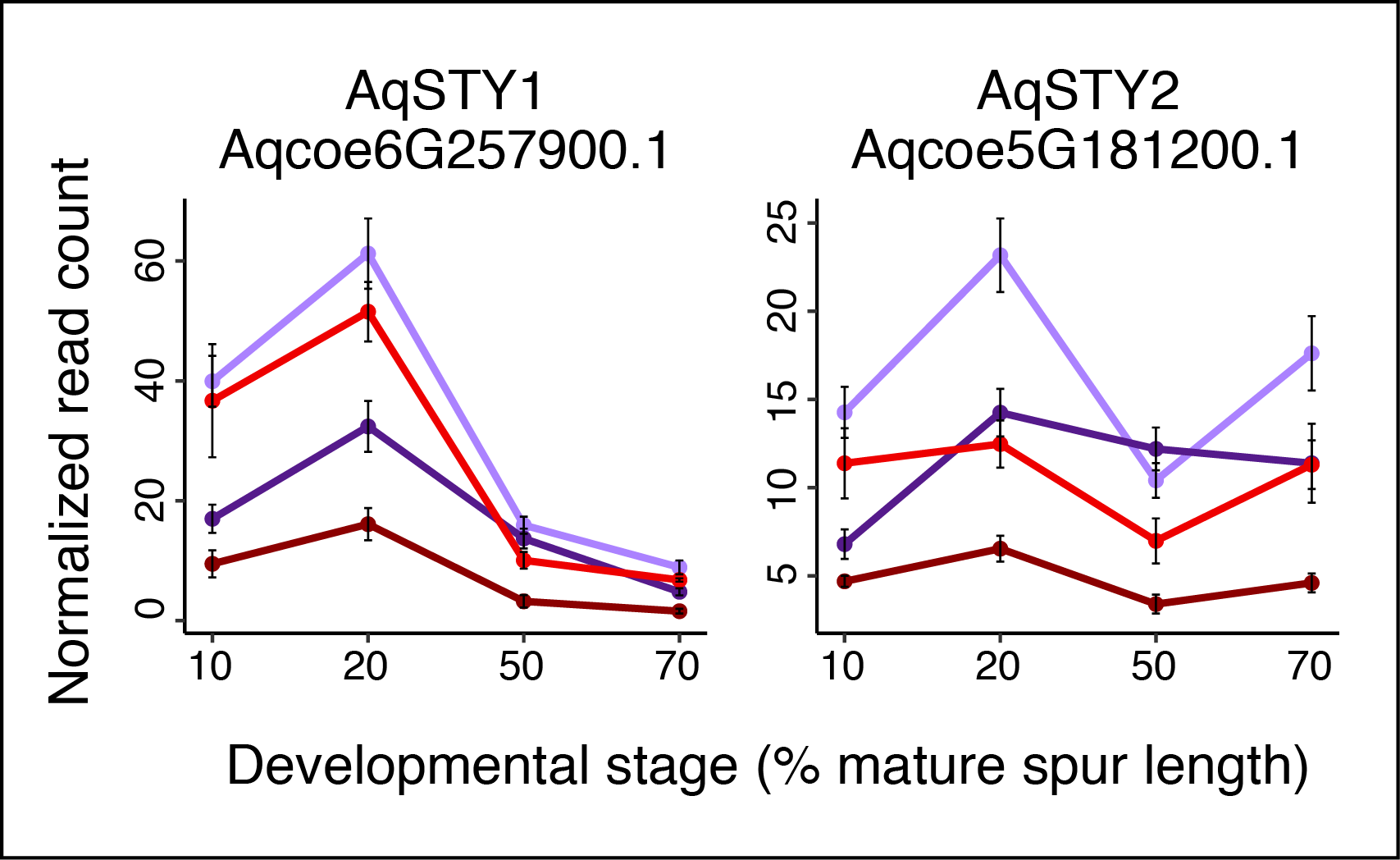

**Appendix S15.** Expression patterns of *AqSTY1* and *AqSTY2* by species, tissue, and developmental stage. Mean normalized read counts across all biological replicates are shown, with error bars representing the standard error of the mean. Light purple, *A. brevistyla* proximal tissue; dark purple, *A. brevistyla* distal tissue; light red, *A. canadensis* proximal tissue; dark red, *A. canadensis* distal tissue.
